# A dendritic cell population responsible for transglutaminase 2-mediated gluten antigen presentation in celiac disease

**DOI:** 10.1101/2025.06.02.657527

**Authors:** Fu-Chen Yang, Harrison A. Besser, Hye Rin Chun, Megan Albertelli, Nielsen Q. Fernandez-Becker, Bana Jabri, Chaitan Khosla

## Abstract

In celiac disease (CeD), a gluten-dependent autoimmune disorder, transglutaminase 2 (TG2) deamidates selected glutamine residues in gluten peptides, while HLA-DQ2 presents deamidated antigens to inflammatory T cells. The cellular sources of pathogenic TG2 and DQ2 are unclear. Using chemical biology tools, we show that intestinal CD103^+^ dendritic cells (DCs) couple cell-surface TG2 to the endocytic LRP1 receptor to simultaneously deamidate gluten antigens and concentrate them in lysosomes. In DQ2-transgenic mice, CD103^+^ DCs loaded with deamidated antigens migrate from intestinal lamina propria and Peyer’s patches into mesenteric lymph nodes, where they engage T cells. In turn, gluten antigen presentation upregulates intestinal TG2 activity. The tool (HB-230) used to establish a role of CD103^+^ DCs in gluten antigen presentation and TG2 activation in mice also revealed that the TG2/LRP1 pathway is active in human CD14^+^ monocytes. Within this population of circulating monocytes, a DC subset with the gut-homing β7-integrin marker is elevated in CeD patients with active disease compared to non-celiac controls or patients on a gluten-free diet. Our findings not only inform the cellular basis for gluten toxicity in CeD but they also highlight the immunologic role of an enigmatic protein of growing therapeutic relevance in CeD and other immune disorders.

## Introduction

Celiac disease (CeD) is a chronic autoimmune disorder characterized by persistent inflammation of the small intestine that affects approximately 1 in 100 individuals worldwide. Notwithstanding advances in our understanding of disease pathobiology, the current management of CeD remains limited to a strict lifelong gluten-free diet (GFD). Therefore, identifying pharmacologically relevant mechanisms that underpin the onset and/or persistence of CeD remains a priority.

CeD is triggered in genetically predisposed individuals by the ingestion of gluten, a family of proteins found in wheat, barley, and rye. Partially digested gluten peptides are believed to cross the intestinal epithelium and be taken up by antigen-presenting cells (APCs) in the lamina propria below, where they are presented to gliadin-specific CD4^+^ T cells. Over 90% of individuals with CeD express human leukocyte antigen (HLA)-DQ2.5 (encoded by DQA1*0105 and DQB1*0102; hereafter referred to as HLA-DQ2), while a smaller proportion of patients express HLA-DQ8.

In addition to dietary gluten and HLA, the enzyme transglutaminase 2 (TG2) plays a central role in CeD pathogenesis by deamidating antigenic gluten peptides (Figure 1A), which dramatically enhances their affinity for HLA-DQ2/8. Blocking TG2 activity was shown to prevent gluten-induced tissue damage in the small intestine in a genetically engineered mouse model of CeD (1). More recently, an irreversible TG2 inhibitor, ZED-1227, was reported to prevent villus atrophy and crypt hyperplasia in CeD subjects challenged with daily gluten over a period of 6 weeks (2). While a compelling case can thus be made for a causative role of TG2 activity in CeD pathogenesis, the cellular source(s) of the active TG2 that generates deamidated gluten antigens in the celiac small intestine remains unclear. This question is complicated by the abundance of the TG2 protein in the small intestinal mucosa, both inside and outside cells (3). In the cytosol, TG2 is reversibly maintained in a catalytically inactive state by guanine nucleotide binding (4), whereas outside cells TG2 is inactivated by a different allosteric mechanism, a reversible disulfide bond (5). Recent studies have also implicated luminal TG2 as a possible source of pathogenically relevant activity in the celiac small intestine (6).

**Figure 1.**
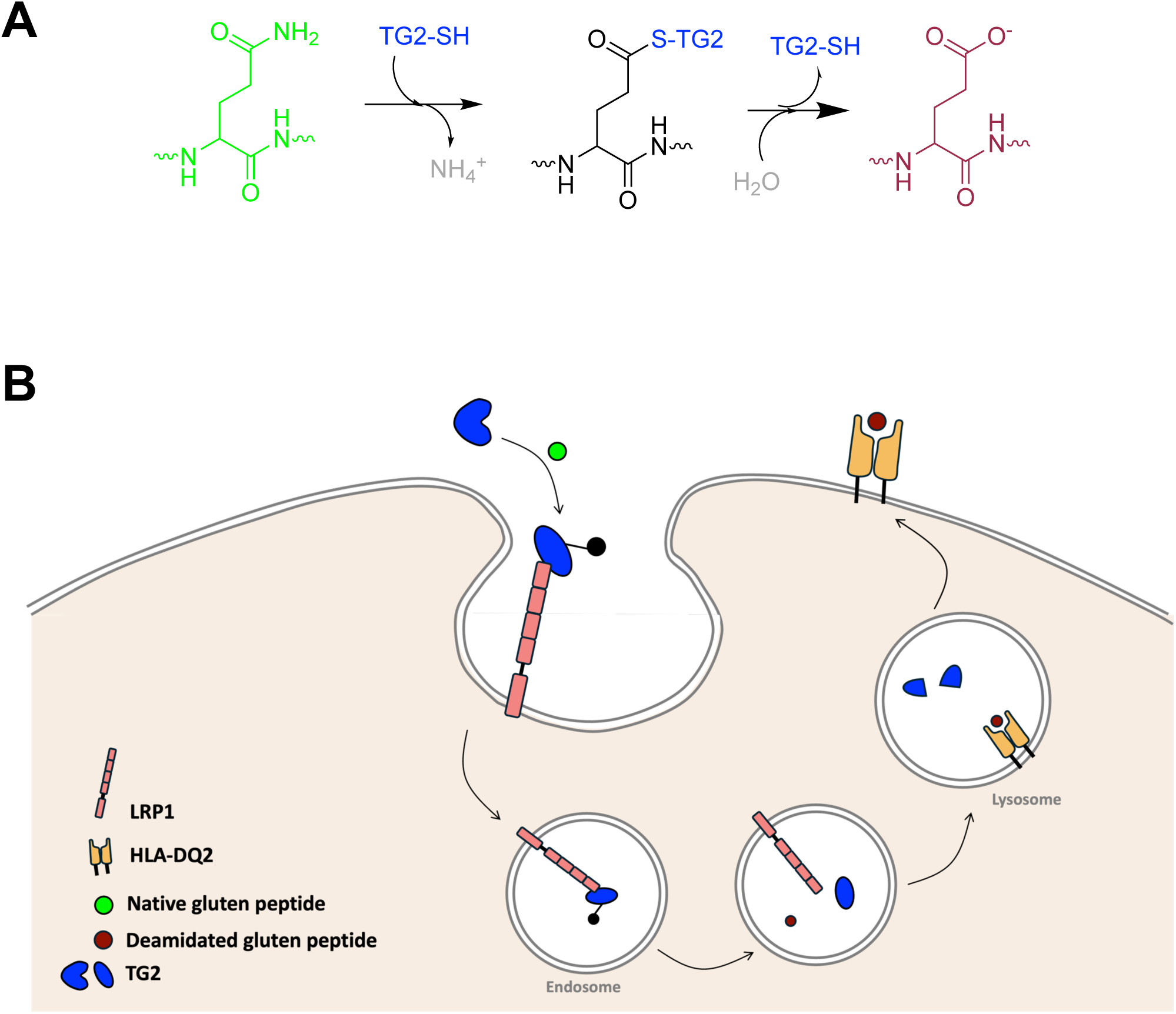
The TG2/LRP1 pathway allows simultaneous receptor mediated endocytosis and deamidation of antigenic gluten peptides. (A) Transglutaminase 2 (TG2) catalyzes deamidation of selected Gln (Q) residues in its substrates via a two-step mechanism. In the first half-reaction, the thiol (SH) functional group of a Cys residue in the active site of TG2 forms a covalent thioester intermediate with the reactive Gln residue, resulting in the release of an ammonium (NH_4_^+^) ion. The second half-reaction involves hydrolysis of this thioester intermediate resulting in a net conversion of the Gln residue into a Glu (E) residue. Alternatively, if a suitable primary amine is available, the thioester intermediate can undergo aminolysis in the second half-reaction, leading to formation of an isopeptide bond (not shown). (B) On the surface of certain antigen-presenting cells, catalytically active TG2 (blue) captures gluten antigens (green) as their corresponding thioester intermediates (black). However, before the thioester intermediate can undergo hydrolysis, it is recognized by the low-density lipoprotein-related protein 1 (LRP1) receptor on these antigen-presenting cells. Binding to LRP1 stabilizes the transient enzyme-substrate complex while activating receptor mediated endocytosis. The TG2-bound thioester is released and hydrolyzed in the acidic environment of endosomes, leading to endosomal delivery of the deamidated gluten peptide (red). Once in the lysosome, the deamidated gluten peptide is loaded into the antigen binding pocket of HLA-DQ2 while the TG2 protein is degraded.

We recently described a hitherto unrecognized mechanism, hereafter referred to as “the TG2/LRP1 pathway”, for gluten antigens to undergo simultaneous deamidation and lysosomal concentration in APCs (Figure 1B) (7). Interaction with low-density lipoprotein-related protein 1 (LRP1) has the remarkable effect of stabilizing a short-lived covalent intermediate between catalytically active TG2 and a suitable substrate for long enough to elicit canonical LRP1 receptor-mediated endocytosis while only allowing hydrolytic resolution of this covalent intermediate in the rapidly acidifying endosome. Guided by the hypothesis that the TG2/LRP1 pathway plays a major role in antigen presentation to HLA-DQ2 restricted, inflammatory T cells in a CeD patient, the goals of our present study were to: (i) identify cells with TG2/LRP1 pathway activity in the small intestine and the periphery; (ii) determine spatial and temporal relationships between TG2 activity associated with the cellular TG2/LRP1 pathway and that which has been previously observed in the extracellular matrix (ECM) of the small intestine; and (iii) demonstrate that at least a subset of these cells can capture dietary gluten antigens and present them to HLA-DQ2 restricted T cells (1, 8). A key finding of our study is that the dominant cells that harness the TG2/LRP1 pathway in the small intestine for gluten antigen presentation are identical to a population of dendritic cells (DCs) previously demonstrated to play a central role in the establishment of dietary antigen tolerance in mice (9, 10). They also show strong similarity to a DC population in the celiac small intestine that undergoes a marked increase in response to dietary gluten and can robustly activate gluten-specific T cells found exclusively in CeD patients (11, 12).

## Results

### Oral HB-230 enables discrimination between ECM-associated and LRP1-associated TG2 activity in the mouse small intestine

Prior to this report, virtually all investigations involving visualization of TG2 activity in the intact mammalian intestine utilized 5-biotinamidopentylamine (5-BP) as a probe (13). Because 5-BP is a co-substrate for the TG2-catalyzed transamidation reaction, it biotinylates proteins with Q (Gln) residues for which TG2 has high substrate specificity. Furthermore, because extracellular TG2 in the small intestine is rapidly deactivated via disulfide bond formation (8), only ECM proteins in the vicinity of the site of TG2 secretion are labeled with 5-BP. Thus, *in situ* 5-BP labeling of intestinal mucosa serves as a convenient marker of TG2 activity, an inference that is underscored by strongly overlapping fluorescence signals from streptavidin and anti-TG2 antibodies in immunohistochemical analysis of intestinal tissues (1, 14).

We recently engineered HB-230, a highly selective, irreversible inhibitor of catalytically active TG2 that also contains sulfocyanine 5 (sulfo-Cy5), a bright fluorophore with maximal absorption at 662 nm (7). The selectivity of HB-230 for TG2 is evident from the large, dose-dependent fluorescent puncta observed in bone marrow-derived dendritic cells (BMDCs) from wild-type (WT) but not TG2-knockout (TG2KO) mice (Figure 2A). The requirement of LRP1 activity for puncta formation is underscored by their strong inhibition by receptor-associated protein (RAP), a selective inhibitor of LRP1-mediated endocytosis (Figure 2B). To visualize the source of catalytically active TG2 that recognizes HB-230, we pre-treated WT BMDCs with HB-230 followed by co-staining with an anti-mouse TG2 antibody. Under these conditions, puncta were observed to be initiated on the cell surface followed by rapid endocytosis and post-endocytic destruction of TG2, as would be expected for most proteins that are trafficked into the lysosome (Figure 2C). Two additional features of HB-230 enabling its oral efficacy as a probe of TG2 activity in the small intestine are its water solubility and acid stability.

**Figure 2.**
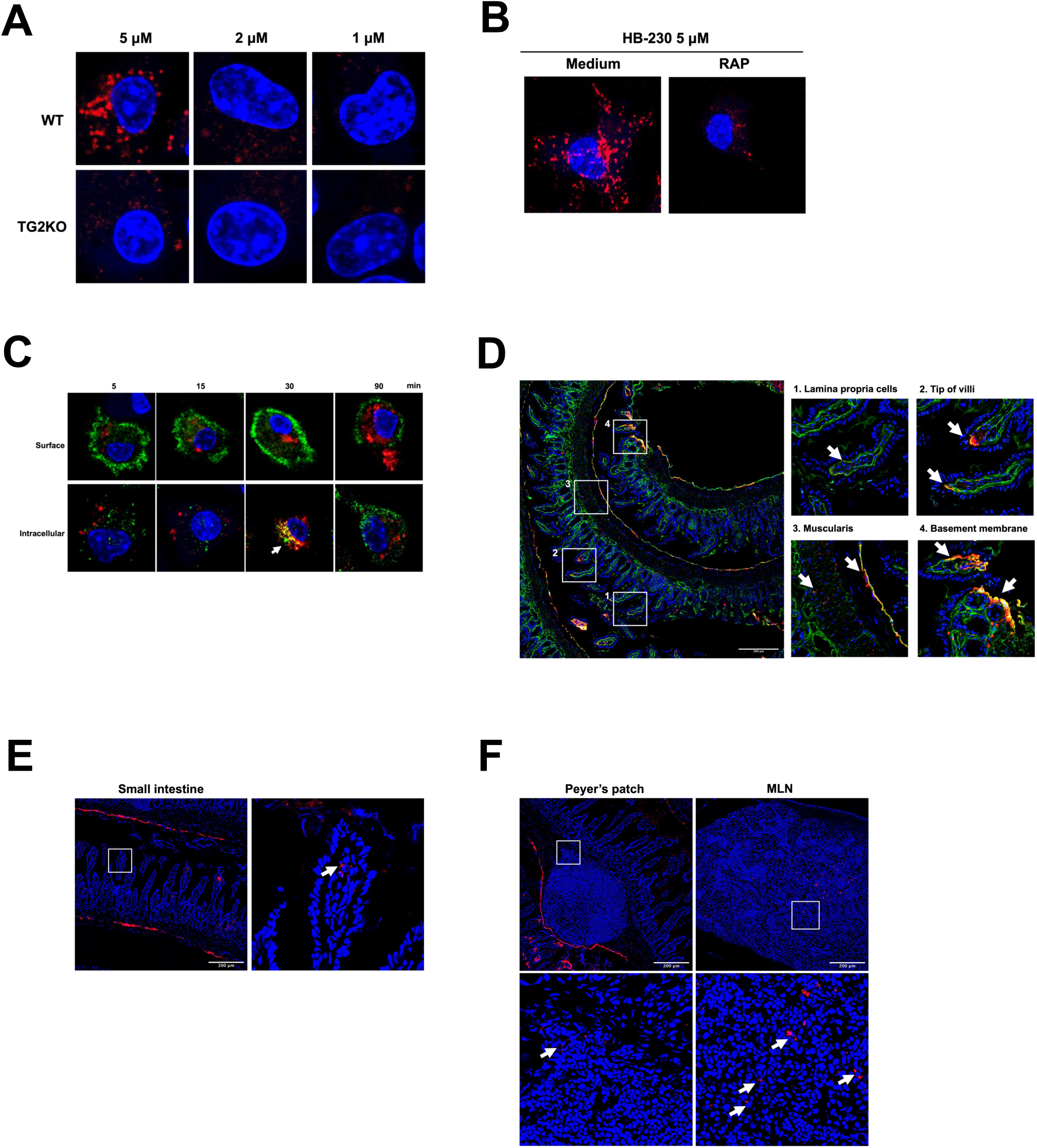
Detection of different active forms of TG2 by HB-230. (A) Bone marrow-derived dendritic cells (BMDCs) from wild-type (WT) and TG2 knockout (TG2KO) mice were treated with increasing concentrations of HB-230 for 90 min. WT BMDCs showed prominent HB-230 puncta, corresponding to lysosomes, which were absent in TG2KO cells. (B) WT BMDCs pretreated with the LRP1 antagonist RAP (3 μM, 30 min) exhibited reduced HB-230 puncta formation following 90 min of probe treatment, indicating LRP1 is required for lysosomal delivery of TG2. (C) WT BMDCs were incubated with HB-230 (2 μM) for 5–90 min and co-stained with anti-TG2 antibody (green) and DAPI (blue), with or without cell permeabilization to detect surface or intracellular TG2, respectively. Early time points showed low red:green signal at the surface, suggesting limited catalytic activity of TG2. In permeabilized cells, HB-230 and TG2 co-localized at intermediate time points, consistent with lysosomal processing of TG2. (D) WT mice were orally dosed with HB-230 (10 mg/kg), and tissues were collected 2 h later. Ileal cryosections showed four HB-230 labeling patterns: (1) lysosomal puncta in lamina propria, (2) subepithelial extracellular matrix (ECM) co-localization with TG2 at villus tips, (3) sporadic ECM labeling in the muscularis, and (4) basement membrane of villi. (E, F) Additional HB-230-labeled cells were observed in (E) the lamina propria and (F) Peyer’s patches and mesenteric lymph nodes (MLN), with perinuclear puncta (white arrows) suggestive of lysosomal localization. All images were acquired at 40× magnification.

As a step toward studying the TG2/LRP1 pathway in the mouse small intestine, WT mice were given 10 mg/kg HB-230 orally after fasting overnight (∼16 h). Two hours after oral gavage of HB-230, the small intestine was removed and processed using Swiss-roll tissue preparation methods prior to snap-freezing. From this Swiss-roll preparation, 5-μm cryosections were cut, co-stained with a polyclonal rabbit anti-mouse TG2 antibody and imaged via confocal microscopy. Consistent with our prior observations with 5-BP, sporadic TG2 activity was detected in the ECM of the lamina propria (mostly near the villus tip) and the muscularis of the WT mouse small intestine (Figure 2D boxes 2-4) but not in TG2KO mice (Supplemental Figure 1A). ECM-associated TG2 activity typically appears as a fluorescent yellow signal, presumably due to strong overlap between the red (HB-230) and green (anti-TG2 antibody) fluorophores. Notably, WT (but not TG2KO) mice also contained few sporadic cells with large HB-230 puncta in the lamina propria (LP) cells (Figure 2D box 1). Unlike ECM-associated TG2 activity, this form of punctate HB-230 staining does not overlap with immunohistochemical staining of the TG2 protein; this is consistent with our observation of rapid TG2 degradation by cells with LRP1-associated TG2 activity.

To better understand the dynamics of tissue distribution and localization of HB-230 staining following its oral administration, draining Peyer’s patches and mesenteric lymph nodes (MLNs) were examined from WT and TG2KO mice dosed orally with HB-230. The small intestine containing Peyer’s patches was dissected 2 h post-dose and processed using Swiss-roll tissue preparation before being snap-frozen. MLNs were collected, embedded, and snap-frozen separately. The Swiss-rolled small intestine provided a detailed view of the tissue with the most structure, showcasing cross-sections of villi and Peyer’s patches. The ileum was centered, with the spiral leading outward into the distal jejunum (as shown in Supplemental Figure 1A, left panel). HB-230 puncta (indicative of LRP1-associated TG2 activity) were sporadically distributed throughout the mouse small intestine, from the jejunum to the ileum, with relatively few occurrences in the distal jejunum (Figure 2E, Supplemental Figure 1A). No punctate HB-230 staining was observed in TG2KO tissue, indicating selective labeling of active TG2 by HB-230 (Supplemental Figure 1A). After 2 h of HB-230 administration, small numbers of punctate, HB-230-labeled cells were observed in the villi (Figure 2E), Peyer’s patches and MLNs (Figure 2F) from WT mice. By 24 h, the presence of labeled cells was limited to the MLN (Supplemental Figure 1B). These observations suggested that either the endocytosed fluorophore had been metabolized or otherwise eliminated, or that the labeled cells- potentially migratory DCs- had migrated out of the lamina propria and Peyer’s patches after internalizing the TG2-HB-230 complex.

### CD103^+^ dendritic cells are the predominant source of LRP1-associated TG2 activity in the mouse small intestine

Migratory CD103^+^ DCs are known for sampling dietary antigens in the lamina propria of the small intestine and migrating to draining MLNs where they activate dietary-specific T cells, particularly driving T-helper 1 type immunity to dietary antigens, including gluten, in the presence of inflammatory triggers (9, 10, 15–18). To determine whether CD103^+^ DCs efficiently uptake HB-230 *in vivo*, we co-stained tissue sections with anti-CD103 and anti-CD11c antibodies. Confocal microscopy clearly demonstrated that HB-230^+^ cells were predominantly labeled with an anti-CD103 antibody, and these CD103^+^ cells were co-stained with CD11c (white arrows, Figure 3A). To verify this finding, we isolated lamina propria cells from WT and TG2KO mice dosed orally with HB-230 and subjected them to flow cytometric analysis. After removing dead cells and debris, CD45^+^ cells were enriched using magnetic beads. Focusing on the CD11c^+^MHCII^+^F4/80^-^ population, HB-230 was observed to be taken up most effectively by the CD103^+^ sub-population of DCs in WT mice (Figure 3B). An insignificant fraction of equivalently labeled cells was detected in TG2KO mice (Figure 3B). In contrast to CD103^+^ DCs, MHCII^+^F4/80^+^ macrophages in the small intestine also showed measurable uptake of HB-230, but the difference in labeling frequency between macrophages from WT and TG2KO mice was not significant (Supplemental Figure 2).

**Figure 3.**
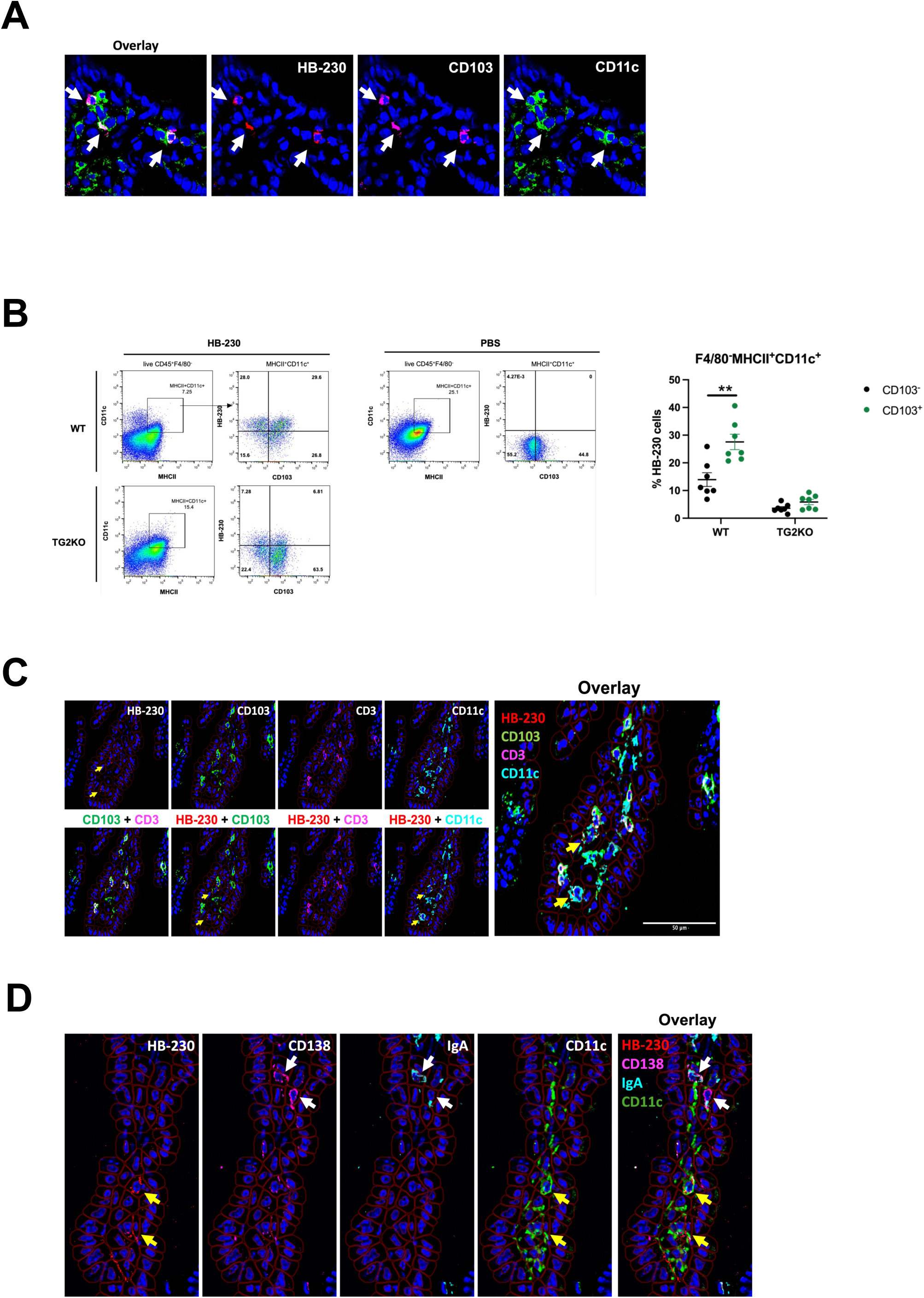
CD103^+^ DCs are predominantly labeled by HB-230 in intestinal lamina propria. Mice received HB-230 at a dose of 10 mg/kg via oral administration. Two hours post-administration, small intestines were collected for either snap freezing or the isolation of CD45^+^ cells from the lamina propria. (A) WT ileum cryosections were co-stained with anti-CD103 and anti-CD11c antibodies. White arrows indicate punctate HB-230 labeled cells co-stained with DC markers. Representative images were captured under a confocal microscope at 40x magnification. (B) Viable CD45^+^ immune cells were enriched from the small intestines of HB-230-dosed WT and TG2KO mice. DCs were gated by MHCII^+^CD11c^+^F4/80^-^ cells and the percentage of HB-230 was analyzed based on PBS control (WT, n=7 and TG2KO, n=7). Small intestines from HB-230-administered WT mice were co-stained with various antibodies (listed in Table 1) using the PhenoCycler multiplex staining system for immune cell characterization. (C) To verify that HB-230^+^CD103^+^ double-positive cells were DCs and not T lymphocytes, tissue sections were additionally stained with an anti-CD3 antibody. Yellow arrows show that HB-230 puncta colocalizes with CD103^+^CD11c^+^ DCs but not with CD3^+^CD103^+^ lymphocytes. (D) To verify that plasma cells did not take up HB-230 to form puncta, tissue sections were additionally stained with an anti-CD138 antibody. Yellow arrows denote HB-230 puncta colocalized with CD11c^+^ DCs but not with CD138^+^IgA^+^ plasma cells (indicated by white arrows). P values were determined using an unpaired Mann-Whitney two-tailed *t*-test. *, *p* < 0.05 and ***, *p* < 0.001. Data represent mean ± SEM.

**Table 1.** Antibodies used in this study.

| **PhenoCycler** |  |  |  |
| --- | --- | --- | --- |
| **Product** | **Clone** | **Barcode** | **Supplier (catalog number)** |
| B220 |  | BX010 | Akoya Biosciences |
| CD103 | 2E7 | BX005 | Thermo Fisher (14-1031-85) |
| CD11b | M1/70 | BX025 | Akoya Biosciences |
| CD11c | N418 | BX022 | Leinco Technologies (C516) |
| CD172a | P84 | BX013 | BioLegend (144002) |
| CD3 |  | BX021 | Akoya Biosciences |
| CD31 | MEC13.3 | BX002 | Akoya Biosciences |
| CD4 | RM4-5 | BX026 | Akoya Biosciences |
| CD45 | 30-F11 | BX007 | Akoya Biosciences |
| CD64 | X54-5/7.1 | BX028 | BioLegend (138302) |
| CD8 | 53-6.7 | BX029 | Akoya Biosciences |
| Collagen A1 | E8F4L | BX001 | Cell Signaling Technology (81375SF) |
| E-cadherin | 24E10 | BX043 | Cell Signaling Technology (96743SF) |
| F4/80 | BM8 | BX020 | Invitrogen (14-4801-85) |
| MHC-II (I-A/I-E) | M5/114.15.2 | BX014 | Akoya Biosciences |
| Ly6C | HK1.4 | BX037 | BioLegend (128002) |
| LYVE-1 | ALY7 | BX032 | Thermo Fisher (14-0443-82) |
| αSMA | | BX055 | Abcam (ab5694) |
| TG2 | House-made | BX019 | Pacific Immunology |
| XCR1 | ZET | BX047 | BioLegend (148202) |
| IgA |  | BX016 | Southern Biotech (1040-01) |
| CD138 | 281-2 | BX004 | BD (553712) |
| **Flow cytometry** |  |  |  |
| **Product** | **Clone** | **Fluorophore** | **Supplier (catalog number)** |
| CD11b | M1/70 | BV421 | BioLegend (101235) |
| MHC II (I-A/I-E) | M5/114.15.2 | FITC | BioLegend (107605) |
| CD11c | N418 | PE | BioLegend (117307) |
| CD103 | 2E7 | PE-Cy7 | BioLegend (121425) |
| F4/80 | BM8 | PerCP-Cy5.5 | BioLegend (123128) |
| XCR1 | ZET | Brilliant Violet 650 | BioLegend (148220) |
| CD14 | HCD14 | FITC | BioLegend (325603) |
| CD11c | 3.9 | PE-Cy7 | BioLegend (301607) |
| CD4 | SK3 | PerCP-Cy5.5 | BioLegend (566923) |
| ITGB7 | FIB504 | PE | BioLegend (321204) |
| **Immunohistochemistry** |  |  |  |
| **Product** | **Clone** | **Fluorophore** | **Supplier (catalog number)** |
| CD11c | HL3 | Biotinylated | BD (5530800) |
| CD103 | 2E7 | Alexa Fluor 594 | BioLegend (121428) |
| CD11b | M1/70 | Purified | Cell Signaling Technology (46512) |
| HLA-DQ2 | SPV-L3 | Alexa Fluor 594 | Biotium (BNC940200-100) |
| Streptavidin |  | Alexa Fluor 488 | Jackson Immuno (016-540-084) |
| Goat anti-rat IgG |  | Cy3 | Abcam (AB98416) |

To assess whether the TG2/LRP1 pathway was active in other types of cells in the lamina propria, we utilized multiplex immunohistochemistry staining of tissues from HB-230 dosed mice. Our 22-plex panel includes lymphocytes (plasma cells, B cells, and T cells) in addition to other relevant cell types (epithelial cells, endothelial cells, lymphatic endothelial cells, and fibroblasts) as well as extracellular matrix components (TG2 protein, collagen 1A1). Intraepithelial lymphocytes have been found to express CD103 (αEβ7 integrin) to mediate T cell adhesion to epithelial cells through its binding to E-cadherin. To investigate whether HB-230^+^ puncta appeared in CD103^+^ T lymphocytes, we examined the CD3^+^CD103^+^population. None of the CD3^+^CD103^+^ T cells were found to have HB-230 puncta (Figure 3C). Because TG2-specific plasma cells have also been observed in substantial numbers in small intestinal biopsies of CeD patients (19, 20), we also sought to test whether the plasma cells present in our samples had an active TG2/LRP1 pathway in the absence of inflammation. For this, tissue samples from WT mice were co-stained with CD138 and IgA to identify IgA-producing plasma cells within the lamina propria. No HB-230 labeling was detected in CD138^+^IgA^+^ plasma cells (Figure 3D).

### ECM-associated TG2 activity colocalizes with type I collagen in lamina propria

Upon secretion by *TGM2*-expressing cells, TG2 interacts with extracellular matrix proteins, including fibronectin and collagens (21). Type I collagen is a major ECM protein in the small intestine, where it supports epithelial cell growth (22). To identify possible sources of the ECM-associated TG2 activity shown in Figure 1D, frozen tissue sections of the small intestine from HB-230-dosed WT mice were first co-stained with antibodies against TG2 and type I collagen. As expected, in the lamina propria of these sections (Supplemental Figure 3A) a clear overlap was observed between sporadic HB-230 (red) and TG2 protein (green); this form of active TG2 also co-localized with type I collagen. Analogous ECM-associated HB-230 was observed along the outer wall of the small intestine.

Because fibroblasts are preferentially localized on the sub-epithelial side of the basement membrane in small intestinal villi (23), we co-stained HB-230-dosed small intestinal tissue sections with an antibody against platelet-derived growth factor receptor α (PDGFRα), a well-established fibroblast marker. Indeed, the sporadic HB-230 observed along the basement membrane frequently co-localized with PDGFRα^+^ cells (Supplemental Figure 3B).

Because the *TGM2* gene is expressed by many cells in the small intestine, we sought to inquire whether other cell types might also contribute to the TG2 activity revealed by HB-230. Antibodies against E-cadherin, α-smooth muscle actin (αSMA), lymphatic vessel endothelial hyaluronan receptor-1 (LYVE-1), and CD31 were used to identify epithelial cells, smooth muscle cells or fibroblasts, lymphatic endothelial cells, and blood vessel endothelial cells, respectively. Whereas a very small percentage of the abundant epithelial and endothelial cells showed HB-230 positivity, some αSMA-positive cells were co-localized with HB-230 (Supplemental Figure 3C). These αSMA-positive cells were observed within the villi, crypts and muscularis, and are presumably of a fibroblast origin.

### Gluten antigens are presented to T cells in MLNs by CD103^+^ DCs

To investigate whether lamina propria-derived CD103^+^ DCs with an active TG2/LRP1 pathway are capable of gluten antigen presentation in mesenteric lymph nodes, we used a humanized DR3.DQ2 mouse that harbors the human HLA-DQ2.5 genes while lacking endogenous mouse MHCII genes (24). A cohort of these transgenic mice were maintained initially under gluten-free conditions, followed by a 14-day enhanced gluten diet in which a well-characterized 33-mer gluten peptide harboring multiple HLA-DQ2.5 epitopes (LQLQPFPQPQLPYPQPQLPYPQPQLPYPQPQPF; (25)) was dosed by oral gavage every other day. On the final day of the study, mice were also dosed with 10 mg/kg HB-298, a sulfo-Cy5 conjugated analog of this 33-mer peptide, either 2 h or 24 h prior to sacrifice. Their small intestines, Peyer’s patches, and MLNs were dissected and snap-frozen for imaging.

In contrast to HB-230 uptake by CD103^+^ DCs in the lamina propria, very few HB-298^+^ cells were detected in the small intestinal villi of this cohort of mice (data not shown). Remarkably though, a significant number of HB-298^+^CD103^+^ double-positive cells were observed in the MLNs, and their abundance remained relatively unchanged between 2 h and 24 h (Figure 4A). A few double-positive cells were also observed in Peyer’s patches 2 h post-dosing, but their numbers declined by 24 h (Figure 4B). To establish whether HB-298 persisted in the MLN by virtue of its ability to bind to HLA-DQ2, we co-stained tissues with the SPV-L3 antibody that specifically recognizes HLA-DQ2. This antibody strongly co-localized with both the anti-CD103 antibody and HB-298 in the MLNs and Peyer’s patches (Figure 4A and B). A similar control experiment performed with WT mice revealed far fewer HB-298^+^cells in the MLN (Supplemental Figure 4A). Our data suggests that HLA-DQ2^+^CD103^+^ DCs are exquisitely well-equipped to sample dietary gluten antigens in the small intestine and/or Peyer’s patches and later migrate to the MLNs to enable antigen presentation.

**Figure 4.**
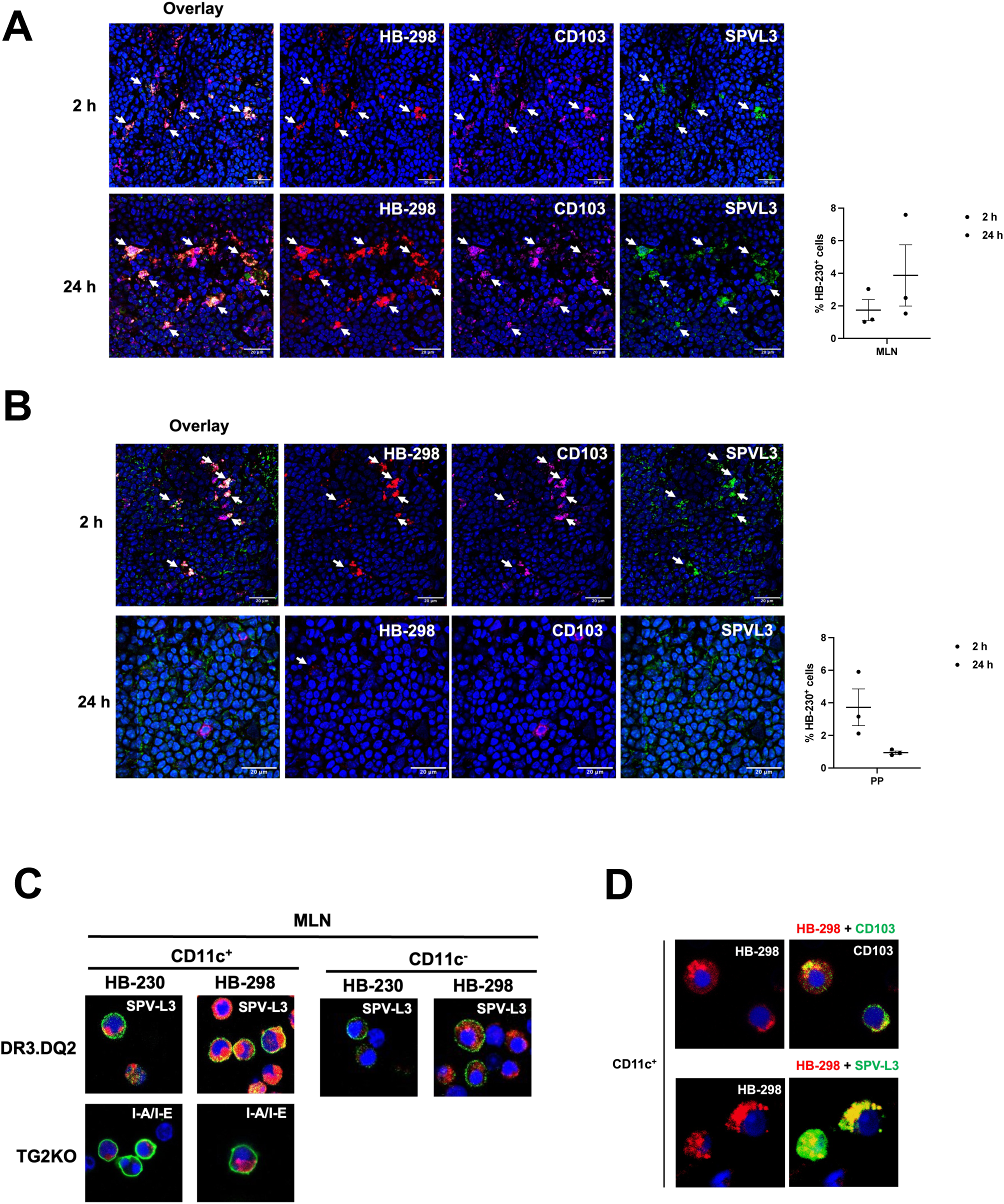
CD103^+^ DCs present gluten peptides in MLN in an HLA-DQ2 restricted manner. DR3.DQ2 mice were orally dosed with 10 mg/kg of HB-298 (a fluorescent analog of an immunodominant 33-mer gluten peptide harboring multiple epitopes recognized by HLA-DQ2) either 2 h or 24 h prior to sacrifice. After euthanasia, mesenteric lymph nodes (MLNs) and Peyer’s patches (PP) were collected, snap-frozen, and stained with antibodies. (A) MLN and (B) PP from DR3.DQ2 mice following HB-298 administration were co-stained with antibodies targeting CD103 and HLA-DQ2 (SPV-L3). White arrows indicate HB-298 labeling of DQ2^+^CD103^+^ DCs. Quantification of labeled cells from MLN and PP confirmed that, whereas the prevalence of HB-298 labeled cells remain unchanged in the MLN at the 24 h time-point, analogously labeled cells in the PP had migrated out of this compartment by 24 h. (C-D) CD11c^+^ and CD11c^-^ cells isolated from the MLNs of DR3.DQ2 and TG2KO mice were incubated with 2 μM of HB-230 or HB-298 for 90 min. (C) Thereafter, MLN cells from DR3.DQ2 mice were stained with antibodies against HLA-DQ2 (SPV-L3), while cells from TG2KO mice were stained with antibodies against mouse MHCII (I-A/I-E). Cell-surface gluten peptide presentation (indicated by a yellow cell boundary) was noted in HB-298-treated CD11c^+^ cells from DR3.DQ2 mice but not in HB-230-treated cells or in TG2KO mice. (D) Also thereafter, HB-298-treated MLN cells from DR3.DQ2 mice were stained with anti-CD103 and anti-DQ2 (SPV-L3) antibodies in CD11c^+^ cells from MLNs of DR3.DQ2 mice. Here too, strong antigen presentation by CD103^+^ DCs was noted. Representative images were captured using a confocal microscope at 40x magnification. Data represent mean ± SEM.

To verify the extraordinary capacity of these dendritic cells to present dietary gluten antigens to T cells, we isolated CD11c^+^ DCs along with CD11c^-^ cells as controls from the MLNs of both DR3.DQ2 and TG2KO mice. These cells were incubated *ex vivo* with either 2 μM HB-230 or HB-298 for 90 min, followed by staining with antibodies against their cognate MHCII molecules. Only HLA-DQ2-expressing CD11c^+^ cells were able to present the fluorescent gluten peptide (but not HB-230) on their cell surface (Figure 4C). Neither CD11c^-^ cells from DR3.DQ2 mice nor CD11c^+^ cells from TG2KO mice showed significant co-localization of HB-298 with their cell-surface MHC II, although peptide uptake was observed in some of these cells. As expected, HB-230 uptake was stronger in double-positive HLA-DQ2^+^CD11c^+^ cells than in either CD11c^-^ cells from DR3.DQ2 mice or CD11c^+^ cells from TG2KO mice, although the fluorophore was not presented on the surface of the double-positive cells. The CD103 positivity of these double-positive DCs was also confirmed (Figure 4D).

### Dietary gluten upregulates ECM-associated and DC-associated TG2 activity in the small intestine of HLA-DQ2 mice

Certain viral infections and other inflammatory triggers have been shown to increase TG2 activity in the small intestine (1, 17). However, the relationship between HLA-DQ2, gluten and TG2, the three principal pathogenically relevant molecules in CeD, is unknown. We therefore sought to test the hypothesis that dietary gluten alone can activate either the cell-associated TG2/LRP1 pathway or ECM-associated TG2 activity (or both) in an HLA-DQ2.5 background.

Parental WT and DR3.DQ2 mice were maintained on a gluten-free diet prior to weaning the pups, and all offspring were maintained on a gluten-free diet until initiation of the study. Our 14-day study involved three cohorts of mice for each genotype – a gluten-free diet, normal chow (containing an indeterminate amount of gluten), and an enhanced gluten diet (described above). After 14 days, an oral gavage of 10 mg/kg of HB-230 was given 2 h prior to sacrifice and harvesting of the small intestines.

No significant differences were observed in small intestinal TG2 activity between WT and DR3.DQ2 mice that were fed either normal chow or a gluten-free diet. In contrast, the enhanced gluten diet led to a marked increase in TG2 activity in DR3.DQ2 but not WT mice (Figure 5A, Supplemental Figure 5A). Gluten-dependent upregulation of TG2 activity was most pronounced in the ECM (Figure 5B), although the prevalence of punctate HB-230^+^ cells in the lamina propria also increased (Supplemental Figure 5B).

**Figure 5.**
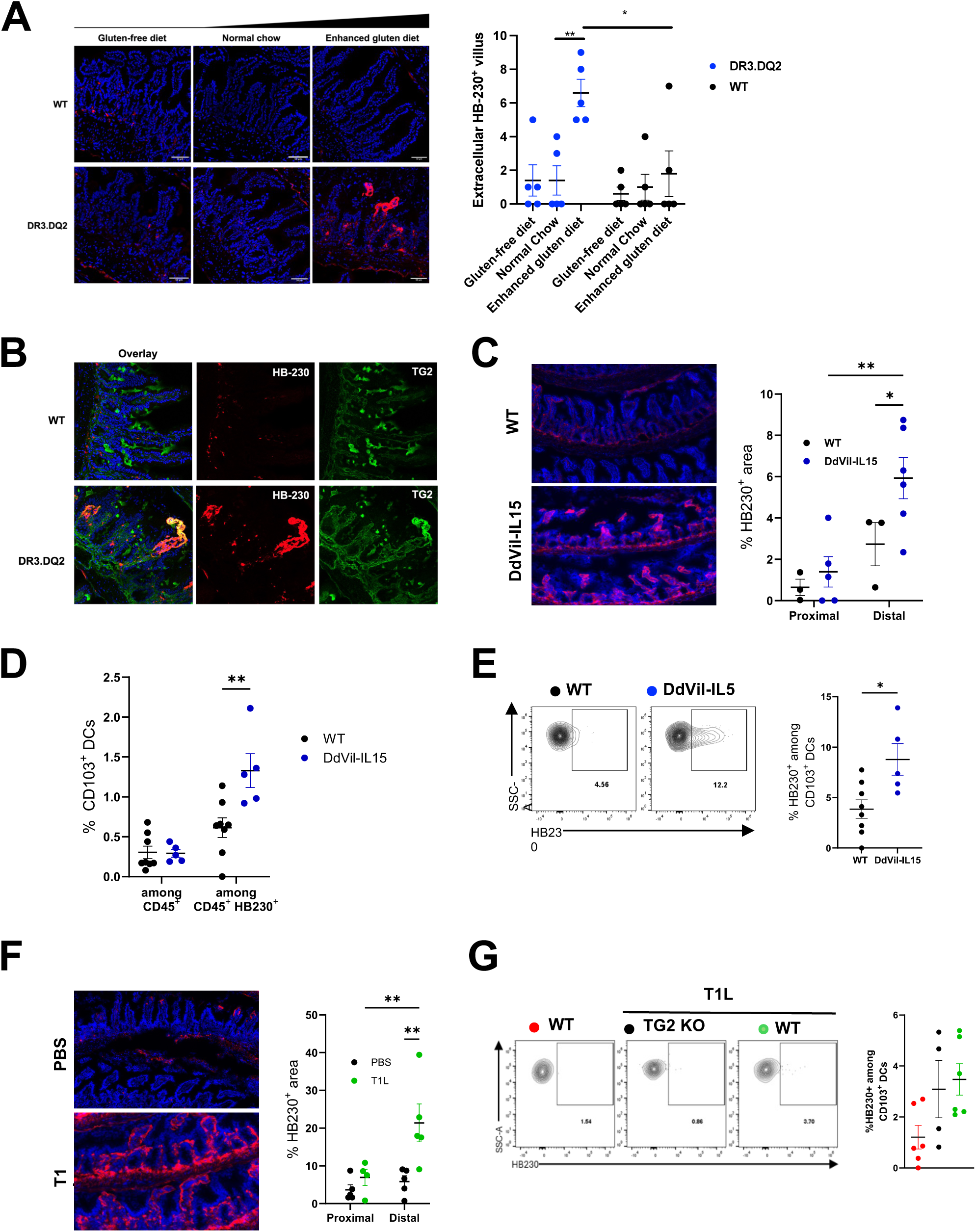
Enhanced gluten diet, reovirus infection, and IL-15 overexpression increase TG2 activity in the small intestine. (A, B) WT and DR3.DQ2 mice were fed control or gluten-enriched diets for 14 days prior to oral HB-230 administration (10 mg/kg). Two hours later, tissues were collected and stained. (A) DR3.DQ2 mice on a gluten-enhanced diet showed increased HB-230 labeling at villus tips. Dot plot quantifies extracellular HB-230⁺ villi per 10 villi. (B) Confocal imaging revealed strong co-localization of HB-230 with TG2 at villus tips, indicating increased ECM-associated TG2 activity. (C–E) DdVil-IL15 transgenic and WT mice received HB-230 (10 mg/kg, oral), and intestines were collected 2 h later. (C) Quantification of HB-230⁺ signal (pixel area relative to total tissue area) in proximal and distal intestine sections revealed increased TG2 activity in DdVil-IL15 mice. (D) Flow cytometry of lamina propria cells showed a higher frequency of CD103⁺ DCs among total CD45⁺ cells and HB-230⁺CD45⁺ cells in DdVil-IL15 mice. (E) Representative flow cytometry plots of HB-230⁺ CD103⁺ DCs in WT (n=8) vs. DdVil-IL15 (n=5) mice. (F, G) WT and TG2 KO mice were infected with T1L reovirus or treated with PBS 46 h prior to HB-230 administration. (F) Confocal images showed increased extracellular HB-230 staining in distal intestines of reovirus-infected WT mice. (G) Percentage and representative plots of HB-230⁺ CD103⁺ DCs in WT (red, n=6), TG2 KO (black, n=4), and T1L-infected WT (green, n=6) groups. Statistical significance: two-way ANOVA with Fisher’s LSD (C, D), unpaired two-tailed t-test (E, F), one-way ANOVA with Tukey’s test (G); *p < 0.05, **p < 0.01. Data shown as mean ± SEM.

### Reovirus infection or IL-15 overexpression also upregulates TG2 activity in the small intestine

IL-15 is an inflammatory cytokine upregulated in reovirus infection; it also plays a critical role in CeD pathogenesis (1, 26). To investigate whether IL-15 directly activates small intestinal TG2, we used DdVil-IL15 transgenic mice, which specifically overexpress IL-15 in their small intestinal epithelium and lamina propria. By quantifying HB-230-positive areas in the small intestine tissue, it was determined that IL-15 overexpression enhanced TG2 activation, particularly in the distal small intestine (Figure 5C, Supplemental Figure 6A).

To analyze the effect of IL-15 overexpression on the prevalence of CD103^+^ DCs or the activity of their TG2/LRP1 pathway, we also performed flow cytometric analysis on isolated lamina propria cells from additional cohorts of DdVil-IL15 and WT mice dosed with HB-230 prior to sacrifice. While the overall DC frequencies do not change significantly between groups, an enrichment of CD11c^+^MHCII^+^ DCs was observed within the CD45^+^HB-230^+^ population in response to IL-15 overexpression (Figure 5D). Notably, DdVil-IL15 mice had more HB-230-positive macrophages and CD103^+^ DCs in their lamina propria compared to their WT counterparts (Figure 5E, Supplemental Figure 7B) but not in lymphocytes (Supplemental Figure 7C). Among all migratory CD103^+^ DCs, CD103^+^CD11b^+^ conventional DC2 had higher TG2/LRP1 activity (Supplemental Figure 7E).

Reovirus infection has been shown to activate TG2 in the small intestine, as assessed with the 5-BP probe (introduced above) (17). To confirm this, T1L reovirus-infected mice and mock-infected controls were dosed orally with HB-230 followed by histologic analysis of Swiss-roll sections (Supplemental Figure 7A). A significant increase in ECM-associated HB-230 was observed in tissue derived from reovirus-infected mice (Figure 5F). TG2 activity was predominantly localized to the distal small intestine, particularly in the ileum and distal jejunum (Supplemental Figure 7B). Notably, CD103^+^CD11b^+^ conventional DC2 had more HB-230 in the lamina propria than in WT or TG2KO mice (Figure 5G, Supplemental Figure 7E, F).

### HB-230 detects differential TG2 activity in peripheral blood monocytes of CeD patients

The above studies demonstrate that a subset of DCs in the small intestinal mucosa of mice are especially well-suited for sampling dietary gluten antigens by virtue of their elevated TG2/LRP1 pathway activity. An earlier analysis of duodenal biopsies from gluten-challenged CeD patients showed an increased number of CD14^+^CD11c^+^ DCs in the lamina propria (11). We therefore sought to assess whether CD14^+^CD11c^+^ DCs in human peripheral blood mononuclear cells (PBMCs) had an active TG2/LRP1 pathway and if so, whether any disease-relevant differences could be observed in this relatively accessible DC population between CeD patients and non-celiac controls. A total of 5 newly diagnosed CeD subjects, 6 CeD subjects on a gluten-free diet, and 3 healthy volunteers were included in this study. PBMCs were incubated with 2 mM HB-230 or PBS for 90 min. After two washes, the cells were stained with an appropriate antibody cocktail and analyzed via flow cytometry. HB-230 positivity was regarded as a marker of cellular TG2/LRP1 pathway activity.

In contrast to CD4^+^ T lymphocytes, for example, which lacked detectable TG2/LRP1 pathway activity, a majority of CD14^+^ cells from both non-celiac controls and CeD patients accumulated HB-230 (Figure 6A). Both CD11c^+^ cells (DCs) and CD11c^-^ cells (monocytes, macrophages) within this population of CD14^+^ cells showed comparable levels of TG2/LRP1 pathway activity (Figure 6B). In healthy subjects as well as CeD patients, a sub-population of CD14^+^ cells had high levels of this marker (CD14^hi^); the TG2/LRP1 pathway was correspondingly more active in these cells (Figure 6C). Thus, cells of the monocytic lineage in human peripheral blood appear to have an active TG2/LRP1 pathway.

**Figure 6.**
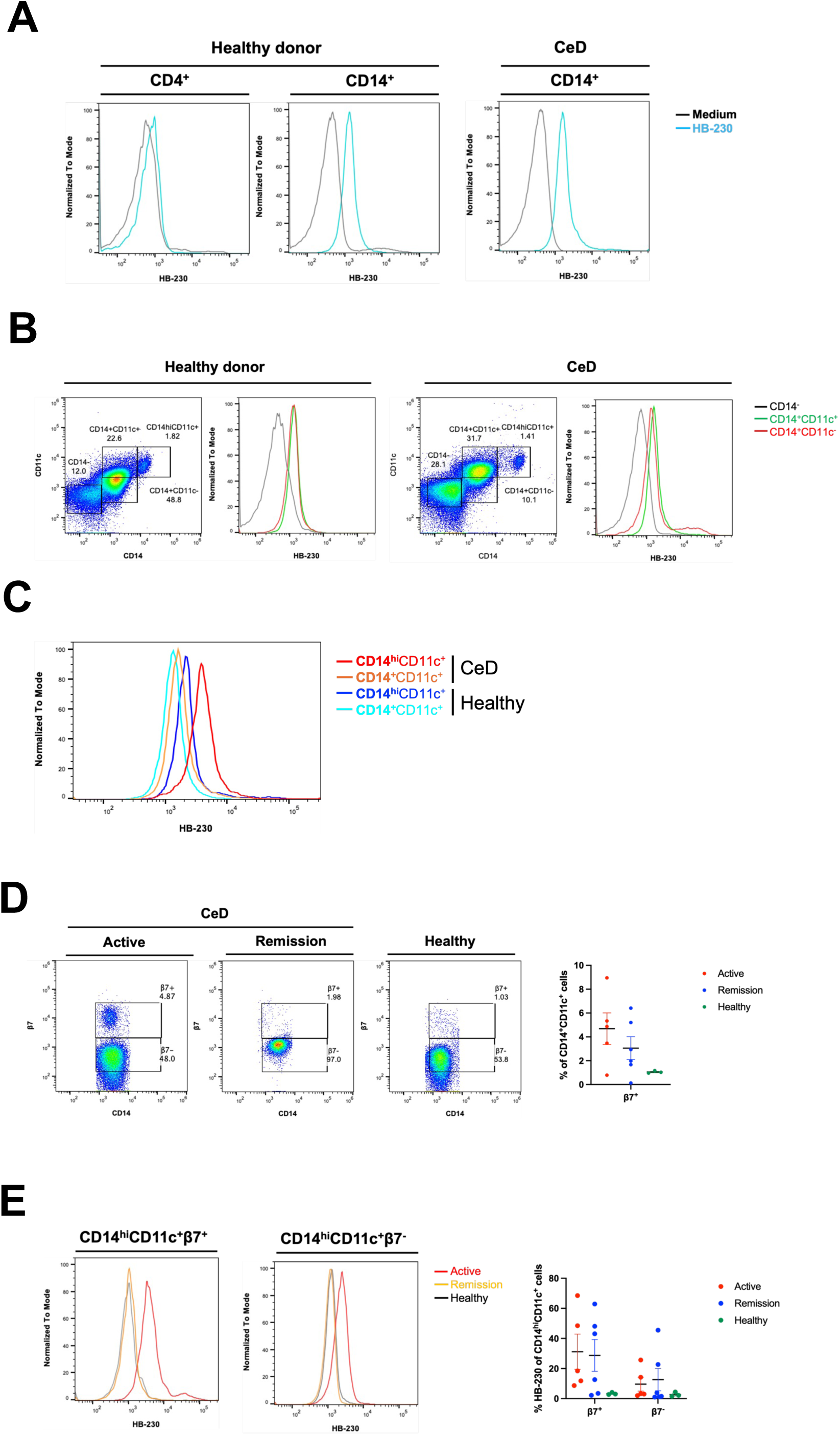
HB-230 uptake by peripheral CD14^+^ monocytes is increased in patients with celiac disease compared to healthy controls. Whole blood from either CeD or healthy subjects was collected. Red blood cells were removed, and the remaining cells were incubated with HB-230 for 90 min. Following incubation, cells were stained with antibodies and analyzed using a flow cytometer. (A) The histogram displays the fluorescence intensity of HB-230 among CD4^+^ T cells or CD14^+^ monocytes from CeD subjects or healthy donors. (B) The dot plot shows the monocyte gating, and the histogram shows the mean fluorescence intensity of HB-230 among CD14^-^ cells (black), CD14^+^CD11c^+^ monocytes (green), and CD14^+^CD11c^-^ monocytes (red) from CeD subjects or healthy donors. (C) The histogram displays the fluorescence intensity of HB-230 among CD14^hi^CD11c^+^ or CD14^+^CD11c^+^ monocytes from CeD subjects or healthy donors. (D) The dot plot displays the percentage of CD14^+^CD11c^+^ monocytes with or without β7-integrin expression in CeD subjects (active CeD, n=5; remission, n=6) or healthy donors (n=3). (E) The histogram displays the fluorescence intensity of HB-230 among CD14^hi^CD11c^+^β7^+^ or CD14^hi^CD11c^+^β7^-^ monocytes from CeD subjects or healthy donors. The dot plot shows the percentage of HB-230 cells among CD14^hi^CD11c^+^β7^+^ or CD14^hi^CD11c^+^β7^-^ monocytes from CeD subjects (active CeD, n=5; remission, n=6) or healthy donors (n=3). Data represent mean ± SEM.

β7-integrin is a marker of gut-associated monocytes and lymphocytes, as it is crucial for adhesion of these migratory cells to MADCAM1 in the small intestine while also aiding in the extravasation of these cells from the bloodstream into gut mucosa. Whereas disease-specific, β7-integrin-positive T cells have been extensively characterized in PBMCs of CeD patients over the past two decades (27–30), to our knowledge an analogous population of gut-homing DCs have not been observed in the peripheral blood of CeD patients. We therefore sought to detect β7-integrin-positive cells within the pool of CD14^+^CD11c^+^ DCs in PBMCs and to assess their TG2/LRP1 pathway activity status. Patients with active CeD revealed a higher prevalence of these gut-homing DCs than either CeD patients on a gluten-free diet or non-celiac controls (Figure 6D). Moreover, most of these cells had an active TG2/LRP1 pathway (Figure 6E). In-depth analysis of these gut-homing DCs with a strong capacity for gluten antigen presentation may shine fundamentally new light on the orchestration of tolerogenic versus inflammatory T cell responses in CeD patients and HLA-matched controls.

## Discussion

For many years now, the central role of the inflammatory T cell response to dietary gluten antigens in celiac disease (CeD) has been widely recognized (28, 31). Two proteins are known to play an essential role in this immune recognition process – HLA-DQ2.5 (or, less frequently, -DQ8), and the enzyme transglutaminase 2 (TG2). Whereas the molecular basis for their association with CeD is well understood (3, 32), their pathogenically relevant cellular origins have remained a mystery. Our findings reported here provide fundamentally new insights into both these questions.

HB-230 is a recently reported small molecule fluorophore that is also a highly selective, active site-directed, irreversible inhibitor of TG2 (7). By developing protocols to use it as a probe of catalytically active TG2 in the small intestine of mice, we have uncovered two principal sources of mucosal TG2 activity. One source of active TG2 is ECM-associated and co-localizes with collagen deposits in the basement membranes that line individual villi; it is likely secreted by fibroblasts that play a key role in fabricating this sub-epithelial structure (23). The other pool of TG2 targeted by HB-230 is localized to CD103^+^ dendritic cells (DCs) in the lamina propria; its punctate intracellular appearance stems from receptor-mediated endocytosis and lysosomal delivery via the action of the TG2/LRP1 pathway (Figure 1). Both sources of TG2 activity are sporadic in a healthy small intestine, comprising a small fraction of the total TG2 protein in this organ.

The ability of active TG2 on the surface of DCs to promote LRP1-mediated endocytosis of gluten antigens represents a highly effective mechanism for gluten antigen presentation to CD4^+^ T cells because it simultaneously catalyzes peptide deamidation while also concentrating the antigen in the lysosomal compartment of this important class of antigen-presenting cells (APCs). Among different subpopulations of intestinal DCs, CD103^+^ DCs are known to be especially important in the recognition of dietary antigens. Indeed, we have previously reported that they represent a major class of antigen-presenting cells that recognize gluten-derived antigens in mice with an appropriate MHC background (24). Using HB-298, a fluorescent 33-residue gluten peptide harboring multiple immunodominant T cell epitopes, we demonstrate that these DCs not only are highly capable of gluten antigen uptake and presentation on HLA-DQ2.5 but that they also migrate to mesenteric lymph nodes where they presumably instruct T cells to assume a tolerogenic or inflammatory phenotype.

Although TG2 is an abundant protein in the ECM of lamina propria in healthy mice, it is predominantly maintained in an inactive state (33). A variety of pro-inflammatory triggers are known to upregulate ECM-associated TG2 activity in lamina propria (1, 8, 17). In active CeD, this pool of TG2 can therefore be expected to contribute significantly to the production of deamidated gluten peptides in the sub-epithelial compartment, thereby enabling antigen presentation by other immune cells (e.g., B cells) (34).

Our findings also demonstrated that both epithelial overexpression of IL-15 and reovirus infection markedly increased HB-230 labeling in the distal small intestine, indicative of ECM-associated TG2 activation. Interestingly, while IL-15 overexpression also enhanced TG2 uptake in migratory DCs and macrophages, reovirus infection, despite robust ECM-associated TG2 activity, did not significantly enhance TG2 internalization in either cell subset. These findings suggest that while ECM-associated TG2 activity can be induced by multiple inflammatory stimuli, only specific environmental contexts promote concurrent activation of the TG2/LRP1 pathway in APCs.

Notably, our findings described in this report prompt reconsideration of HLA-DQ2 mice as models for aspects of CeD. While several laboratories have reported the construction and characterization of this strain of transgenic mice, their inability to mount an immune response to dietary gluten has limited their use in CeD research. Using molecularly defined oral antigens at carefully titrated doses, we show that HLA-DQ2 mice can serve as a good model for the loss of oral tolerance to dietary gluten that is observed in CeD patients.

Last but not least, our analysis of DCs found in peripheral blood of humans has confirmed that some circulating DCs have high TG2/LRP1 pathway activity, presumably allowing them to capture antigens harboring TG2 recognition motifs. This finding speaks to the evolutionary relevance of TG2, an enigmatic protein whose biological role remains uncertain due to the lack of phenotypes associated with TG2-knockout mice (14). Moreover, our discovery that CeD patients have circulating DCs with atypically high TG2/LRP1 pathway activity and the β7-integrin marker opens the door to characterizing a long sought-after APC in this autoimmune-like condition.

The findings reported here prompt several new questions. For example, what is the nature of the catalytically active TG2 localized on the surface of DCs? How does it enable potent activation of endocytosis by the LRP1 receptor in an antigen-dependent manner? Do regulatory CD103^+^ conventional DCs in the small intestine transform into inflammatory DCs in response to signals that lead to a breakdown of oral tolerance in CeD patients, or do inflammatory DCs in patients come from the periphery (11, 12)? How do CD103^+^ DCs upregulate ECM-associated TG2 activity? The tools and approaches described in this study represent powerful starting points for future studies aimed at addressing these questions.

## Materials and methods

### Sex as a biological variable

For human and mouse studies, both male and female subjects were included.

### Animals

Age and sex-matched (8- to 12-week-old) C57BL/6J wild-type (WT) mice were from The Jackson Laboratory (JAX000664) and maintained by in-house breeding. DR3.DQ2 (24) and TG2-knockout (TG2KO) (35) mice were housed in a specific pathogen-free environment at Stanford University under a protocol approved by the Administrative Panel on Laboratory Animal Care (APLAC). DdVil-IL15 transgenic mice (1) were maintained under specific pathogen-free conditions at the University of Chicago.

### Reagents

HB-230 and HB-298 (LQLQPFPQPQLPYPQPQLPYPQPQLPYPQPQPF with the sulfocyanine 5 dye conjugated to the N-terminus) were synthesized using general peptide synthesis protocols previously described (7). HB-230 was dissolved in PBS as a 10 mg/mL solution for oral administration. HB-298 was dissolved in water as a 1 mg/mL solution for oral administration. The intact 33mer gluten peptide cited above was also used as a supplement in an enhanced gluten diet, where it was dissolved in water as a 1 mg/mL solution and administered by oral gavage every other day.

### Tissue collection and Swiss-rolled small intestine preparation

Mice were fasted overnight for 14-16 h before receiving oral gavage dosing. Each mouse received a single bolus dose of HB-230 or HB-298 at 10 mg/kg via oral gavage either 2 h or 24 h before sacrifice. After the oral dosing, the mice were euthanized using carbon dioxide and dissected. The entire small intestine was removed, with fat and mesenteric lymph nodes discarded. The small intestine was cut into two segments. Segment 1 included the duodenum to the distal jejunum, while Segment 2 encompassed the remainder of the jejunum to the ileum. Each segment was cut open lengthwise and flushed with PBS to eliminate luminal residue. The distal jejunum from Segment 1 and the ileum from Segment 2 were positioned centrally and spiraled up. The rolled tissue was embedded in the OCT compound and snap-frozen using dry ice. Mesenteric lymph nodes were collected and embedded separately in OCT compound and snap-frozen using dry ice.

### Bone marrow-derived dendritic cell differentiation, stimulation, and analysis

Bone marrow cells were collected from the femurs and tibias of mice using a 26G needle attached to a 1 mL syringe filled with PBS. The red blood cells were removed using a red blood cell lysis buffer (BioLegend), and the remaining cells were resuspended in RPMI 1640 medium containing 10% fetal bovine serum and 1% antibiotics. These cells were plated in a petri dish at a concentration of 2 million cells per 10 mL of medium and supplemented with 10 ng/mL of GM-CSF for 8 days. On days 3 and 6, 10 mL of fresh GM-CSF-containing medium was added, and the suspension cells were harvested on day 7. Two hundred thousand bone marrow-derived dendritic cells (BMDCs) were plated in a poly-L-lysine pre-coated 24-well glass-bottom dish and incubated with 2 μM HB-230 for 90 min. The cells were rinsed with PBS twice, stained with antibodies, and then fixed with 2% paraformaldehyde for 10 min, followed by staining with 4’,6-diamidino-2-phenylindole (DAPI). Stained cells were visualized and analyzed using a Zeiss LSM 980 confocal microscope (Zeiss) and Fiji ImageJ software.

### Immunohistochemical analysis

A fresh frozen section (5 μm) was fixed in 100% acetone for 10 min at room temperature and blocked with 1.5% bovine serum albumin for 30 min. The sections were stained with the fluorochrome-conjugated antibodies listed in Table 1 and kept in a cold room overnight. Secondary antibody staining was performed, followed by Hoechst 33342 staining for nuclei. The section was mounted in 90% glycerol in PBS and visualized and analyzed using a Zeiss LSM 980 confocal microscope (Zeiss) and Fiji ImageJ software.

### Immunocytochemistry

Bone marrow-derived dendritic cells (BMDCs) were plated on a 0.01% poly-L-lysine precoated glass-bottom culture dish and incubated with 2 μM HB-230 for 5, 15, 30, and 90 minutes. After incubation, the medium was aspirated, and BMDCs were rinsed with PBS before surface TG2 staining. Paraformaldehyde-fixed and Triton X-100-permeabilized BMDCs were incubated with 2% fetal bovine serum to block nonspecific antigens prior to staining intracellular TG2 proteins. Co-localization of HB-230 and TG2 proteins was visualized and analyzed using a Zeiss LSM 980 confocal microscope and Fiji ImageJ software.

### Flow cytometry

After removing Peyer’s patches, the small intestine was cut into small pieces, and lamina propria cells were prepared by removing epithelial cells with 1 mM EDTA for 20 min. The cells were then dissociated in collagenase IV medium (1 mg/mL of collagenase with 0.1 mg/mL of DNase I) for 20 min after rinsing the cell pellet with PBS. Viable immune cells from the lamina propria were purified through two steps using magnetic nanobeads. First, a dead cell removal kit (BioLegend) was used to exclude dead cells, followed by CD45^+^ immune cell enrichment using CD45 nanobeads (BioLegend). To analyze DC subsets in the mouse small intestine, freshly isolated cells were blocked with 2.4G2 (anti-Fc receptor) to prevent non-specific staining and stained with the fluorochrome-conjugated antibodies listed in Table 1. Cells were collected and analyzed using a cell sorter SH800S (Sony Biotechnology) and FlowJo software (BD Biosciences).

### T1L reovirus infection model

Mice were inoculated perorally with purified reovirus diluted in PBS using a 22-gauge round-tipped needle. All T1L reovirus-treated mice were given the same dose of 10^8^ PFU of virus and sacrificed for analysis 48 h post-infection.

### Human peripheral blood mononuclear cells analysis

Blood samples were collected from patients with CeD and non-celiac donors. The red blood cells were removed before use of peripheral blood mononuclear cells (PBMCs). PBMCs from CeD patients (newly diagnosed (n=5) or under gluten-free diet (n=6)) and non-celiac donors (n=3) were incubated in RPMI medium containing 2 μM HB-230 for 90 min. Following incubation, the cells were washed twice with PBS that contained 2% FBS and stained with the antibody cocktail detailed in Table 1. The cells were subsequently collected and analyzed using a cell sorter SH800S (Sony Biotechnology) and FlowJo software (BD Biosciences).

### Statistics

Statistical comparison groups were performed using the Mann-Whitney test, two-way ANOVA with Fisher’s least significant difference (LSD) test, one-way ANOVA with Tukey’s multiple comparison test, or 2-tailed Student’s t test. Statistical analysis was performed with GraphPad Prism 10. P values of less than 0.05 were considered significant.

### Study approval

Mice studies were conducted in accordance with protocols approved by the Administrative Panel on Laboratory Animal Care (APLAC) at Stanford University and the Institutional Biosafety Committee and the Institutional Care and Use Committee of the University of Chicago. Blood samples from CeD patients (Stanford Medicine Outpatient Center in Redwood City) and non-celiac donors (healthy individuals from Stanford Blood Center) were collected with permission from the Stanford Institutional Review Board (Protocol 20362).

## Data availability

Data related to the paper are available from the corresponding author on reasonable request.

## Author contributions

F-CY, CK and BJ designed the research and supervised all investigations. F-CY and HRC performed experiments and analyzed data. HAB provided reagents and intellectual input. CK and F-CY wrote the manuscript. All authors provided critical reviews of the manuscript.

## Acknowledgments

This work was supported by a grant from the NIH (R01-DK-063158) to CK and BJ and by a gift to Stanford University from The Snider Foundation. F-CY was supported by the Stanford Medicine Children’s Health Center through its IBD and Celiac Disease Research Program. HAB was supported by the NIH (F30DK132903) and the Stanford University Medical Scientist Training Program (T32GM007365 and T32GM145402). The authors thank the Wu Tsai Institute Neuroscience Microscopy Service and Cell Sciences Imaging Facility at Stanford University for providing Zeiss confocal microscopes and PhenoCycler (CODEX) service.

---

**Supplemental Figure 1.**
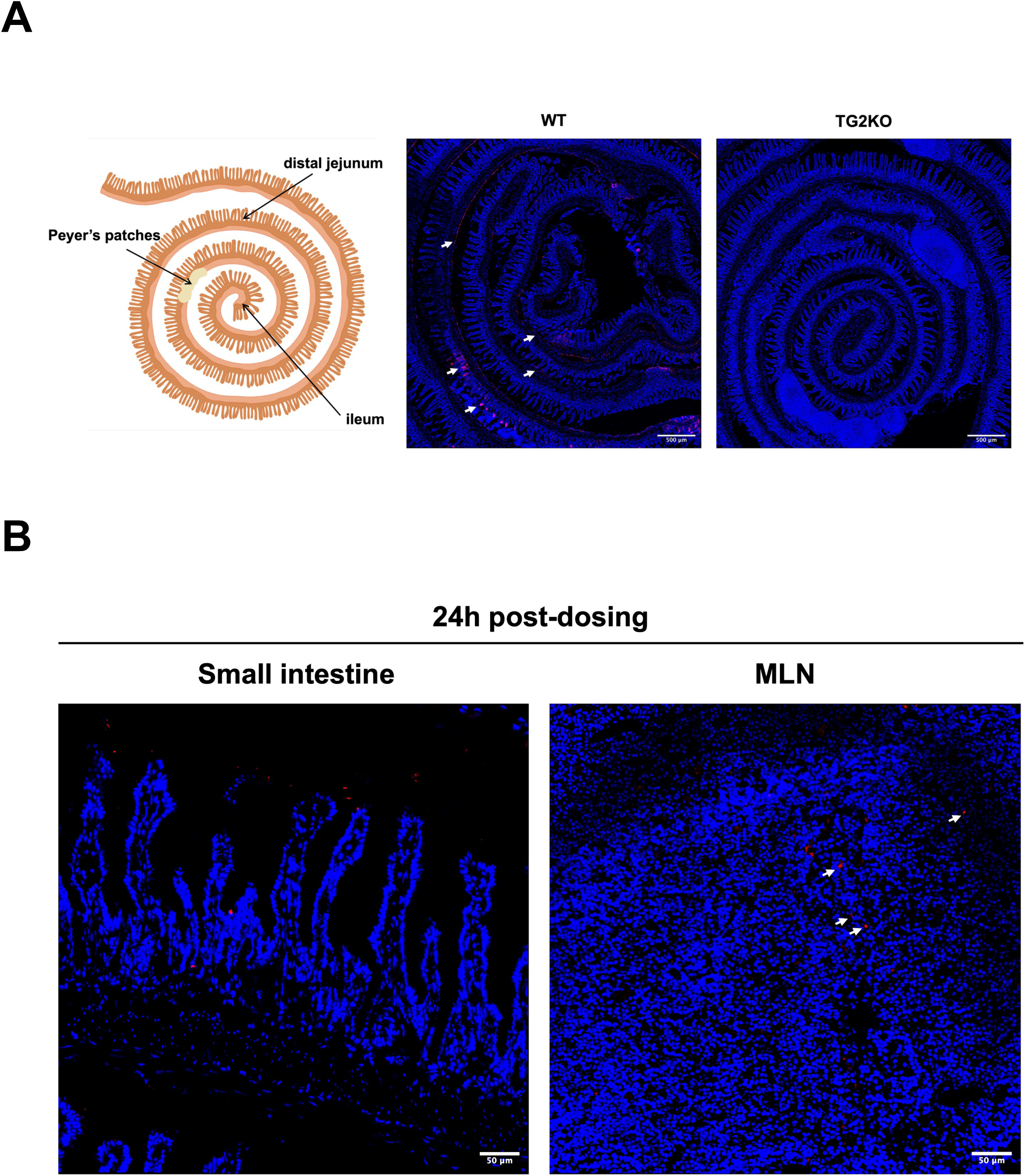
H**B-230-labeled active TG2 is observed in the small intestines of WT mice but not in TG2KO.** (A) A schematic illustration of the Swiss-roll preparation of the ileum, distal jejunum, and Peyer’s patches in the mouse small intestine. Representative images of Swiss-rolls from WT and TG2KO mice dosed with HB-230 2 h before sacrifice. White arrows in the WT image indicate HB-230 signals within the tissue. (B) WT mice received 10 mg/kg HB-230 orally 24 h before euthanasia. After sacrifice, the small intestine and MLN were collected, embedded, and snap-frozen separately. All images were taken using a Zeiss confocal microscope at 40x magnification.

**Supplemental Figure 2.**
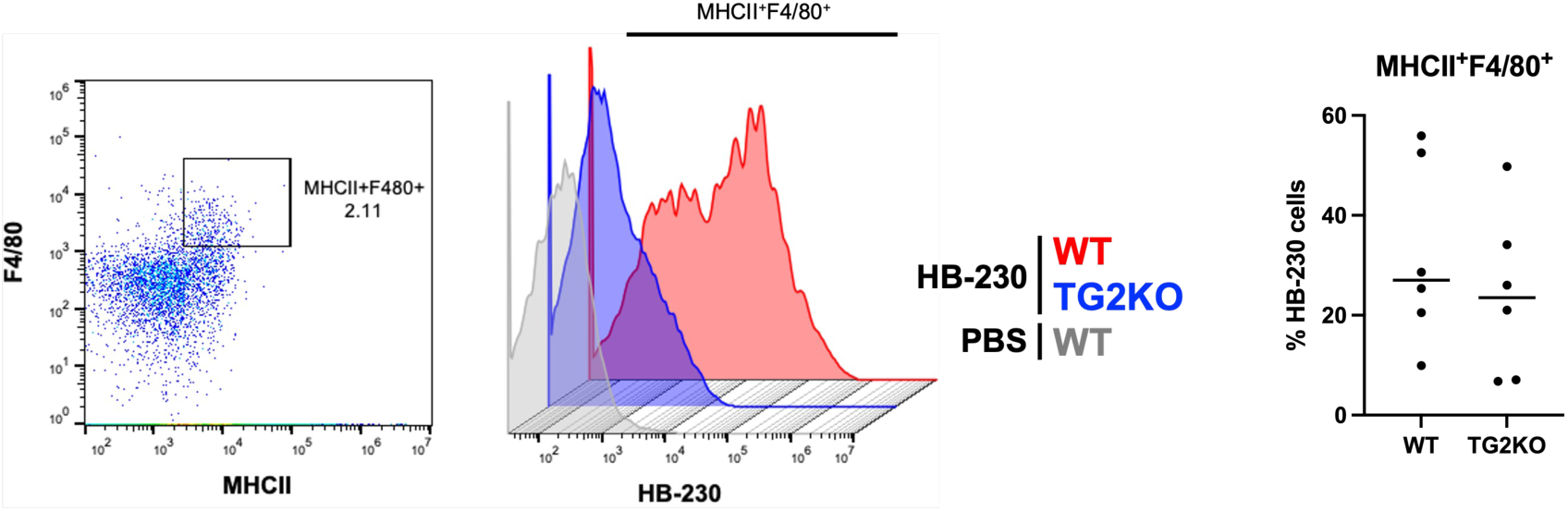
The percentage of HB-230^+^ cells among CD103^+^CD11c^+^MHCII^+^DCs. Small intestinal lamina propria cells were isolated from WT and TG2KO mice after receiving HB-230 2 h before sacrifice. Viable CD45^+^ immune cells were enriched and stained with antibodies. Macrophages were gated on MHCII^+^F4/80^+^ cells. Among macrophages, HB-230 was gated based on the PBS control group (WT, n=7; TG2KO, n=7).

**Supplemental Figure 3.**
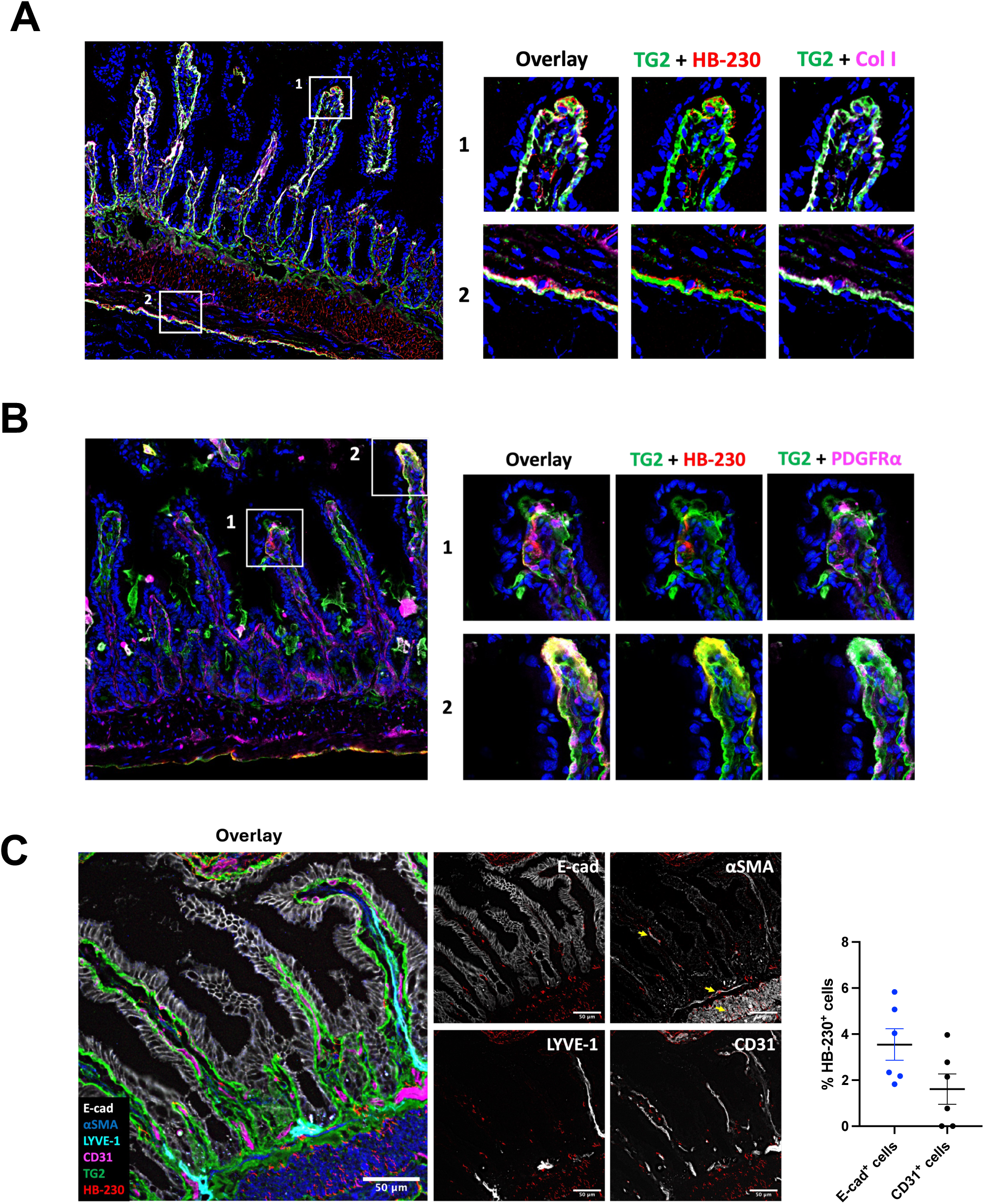
Extracellular HB-230 colocalizes with collagen and smooth muscle actin. WT mice received HB-230 at a dose of 10 mg/kg via oral administration. Two hours after administration, tissues were collected, snap-frozen, and stained with antibodies. Images were captured and analyzed using the PhenoCycler system. (A) (Left) Images of a representative cryosection with extracellular HB-230 overlaid with labeled antibodies against TG2 protein and type I collagen. (Right) Magnified images of the lamina propria (top) and muscularis (bottom). (B) Extracellular HB-230 overlaid with TG2 protein and anti-PDGFRα antibody near the basement membrane of villi. (C) A representative cryosection labeled with HB-230, antibody against TG2 protein, and antibodies against markers of non-immune cell compartments in the small intestine, including epithelial cells (E-cadherin), smooth muscle cells (αSMA), lymphatic vessels (LYVE-1), and endothelial cells (CD31). On the right, % HB-230^+^ cells among epithelial cells and endothelial cells. Each point represents a normalized quantity of E-cad^+^HB-230^+^ cells in E-cad^+^ cells or CD31^+^HB-230^+^ cells in CD31^+^ cells from 10 villi. For each mouse, two randomly picked regions with 10 villi were analyzed (WT, n=3). All representative images were captured using a confocal microscope at 20x magnification. Data represent mean ± SEM.

**Supplemental Figure 4.**
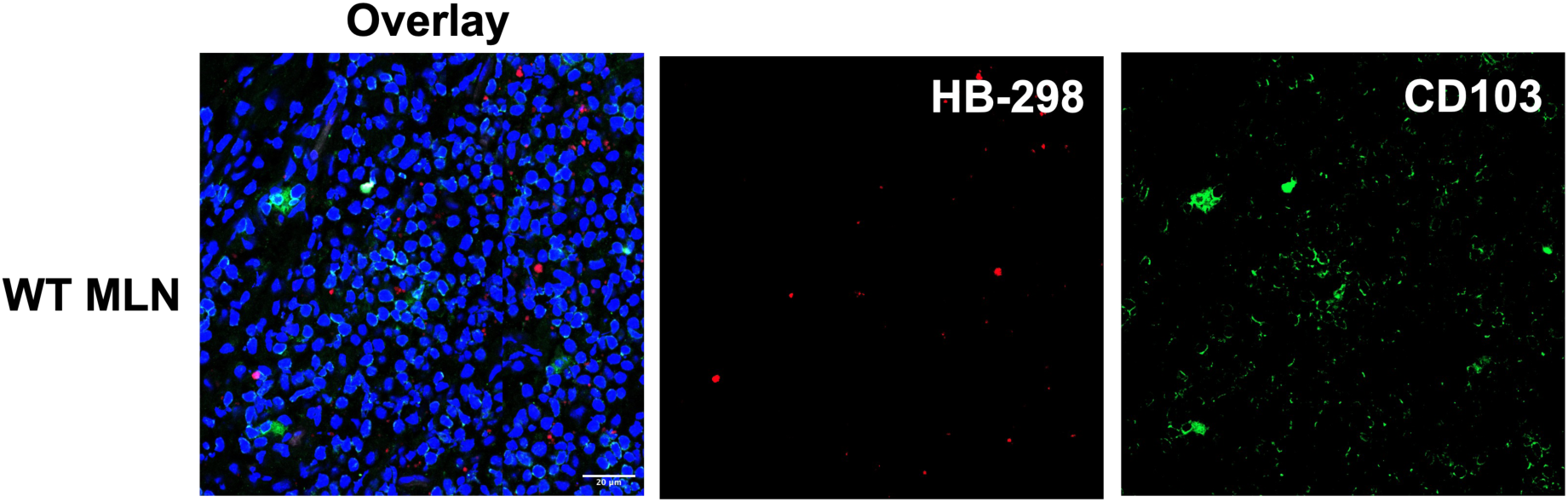
Gluten peptides reach MLNs following oral administration but are not presented by CD103^+^ DCs to T cells. WT mice received 10 mg/kg of HB-298 orally for 2 h before sacrifice. After euthanasia, MLNs were collected, snap-frozen, and stained with anti-CD103 antibody. Some HB-298 is observed in the MLN post-doing but does not colocalize with CD103^+^ DCs. The image was taken using a confocal microscope at 40x magnification.

**Supplemental Figure 5.**
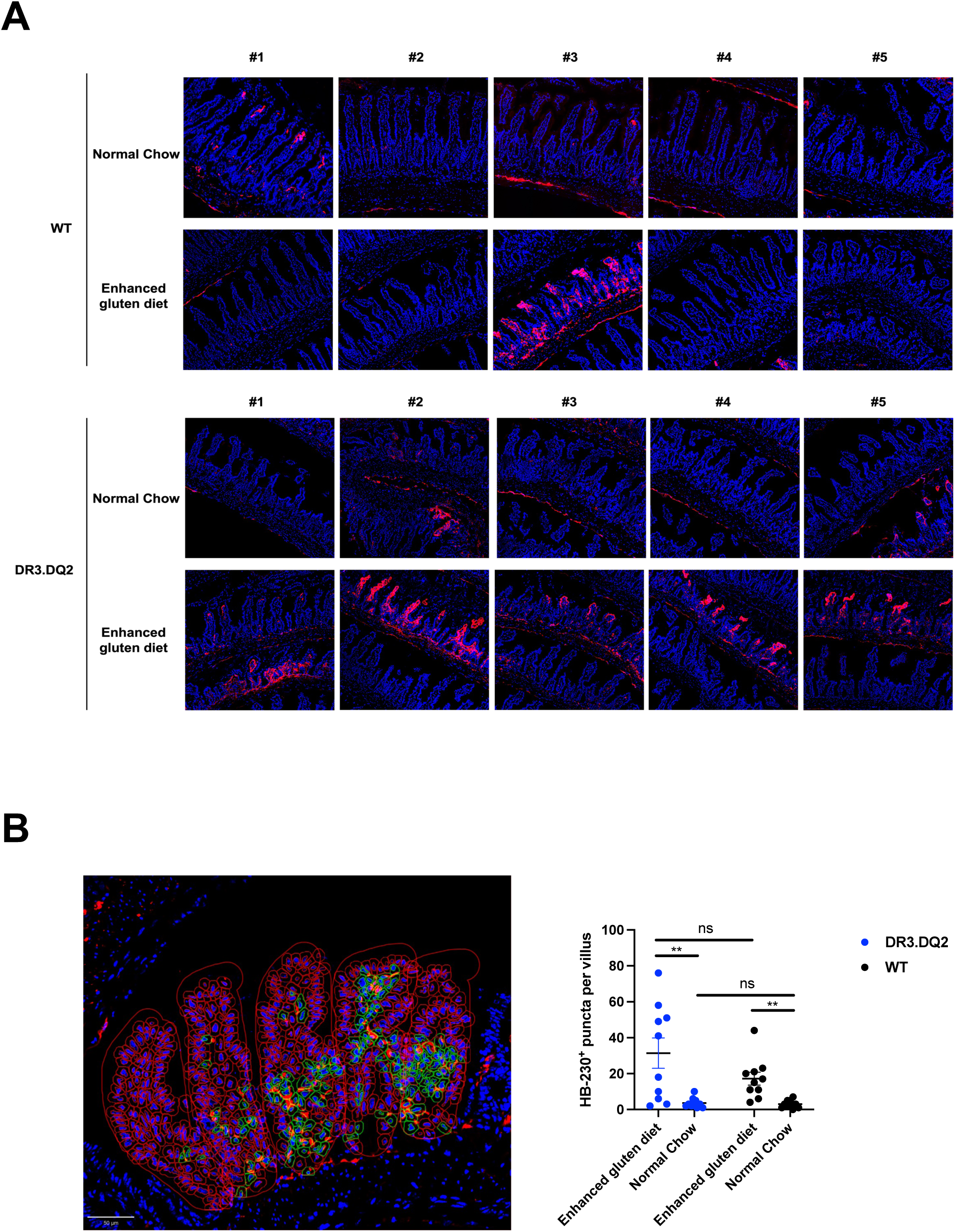
Elevated extracellular HB-230 is observed in DR3.DQ2 mice when on an enhanced gluten diet. WT and DR3.DQ2 mice were given normal chow or specified diets for 14 days prior to receiving 10 mg/kg HB-230 orally. Two hours after administration, tissues were collected, snap-frozen, and stained with antibodies. (A) Elevated extracellular HB-230 was noted at the tips of the villi in DR3.DQ2 mice that received a high-gluten diet for 14 days. Five representative images from 2 to 3 mice were taken using a confocal microscope at 40x magnification. (B) The left image demonstrates that intracellular HB-230 counting using QuPath software (36) to identify HB-230^+^ cells in each villus. The dot plot shows HB-230^+^ cells per villus in each cohort (n=10). P values were determined using an unpaired Mann-Whitney two-tailed *t*-test. **, *p* < 0.01. Data represent mean ± SEM.

**Supplemental Figure 6.**
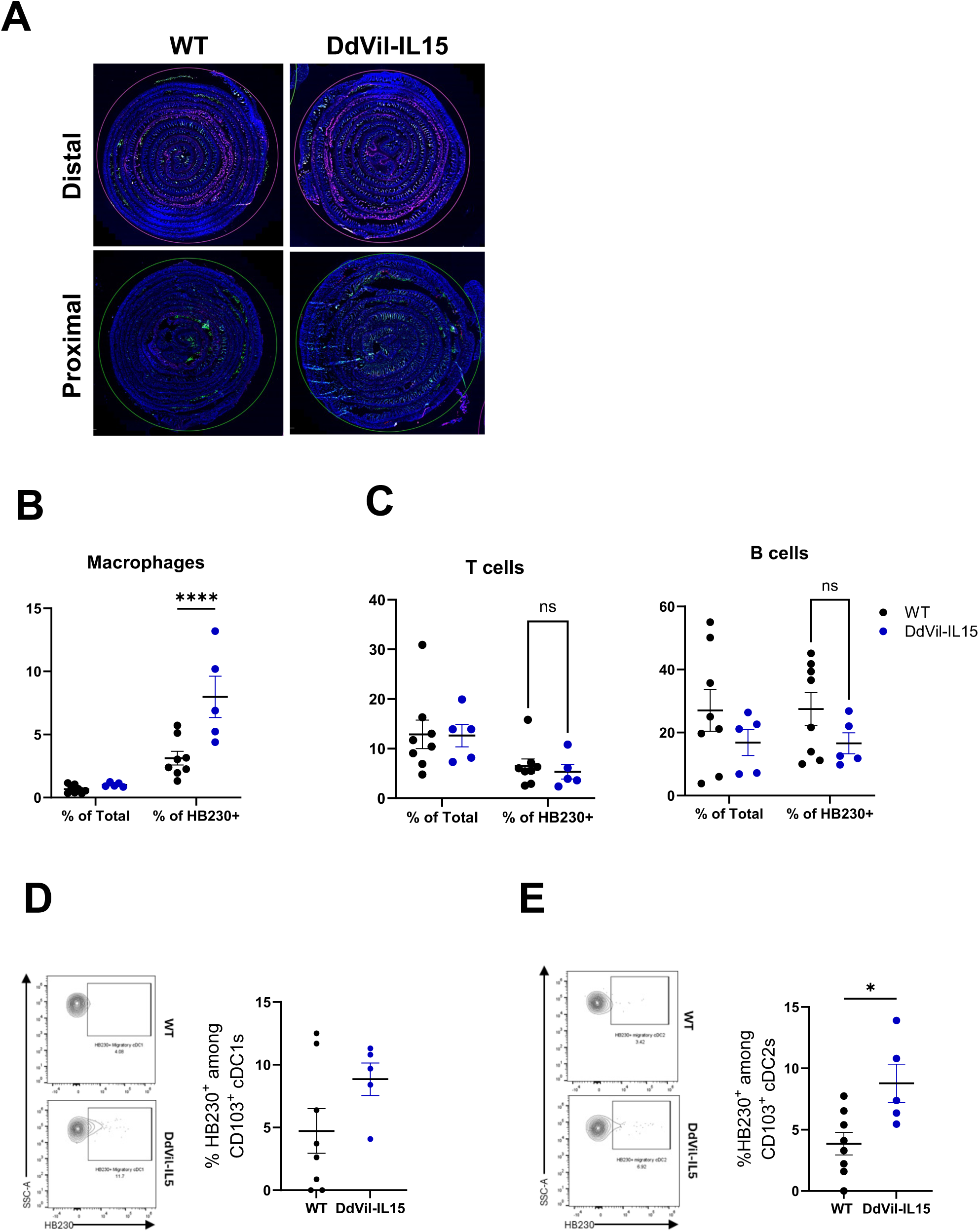
Increased HB-230^+^ cells in macrophages but not T and B lymphocytes in DdVil-IL15 mice. (A) Representative confocal microscopic images of distal and proximal Swiss-rolled small intestine sections of WT or DdVil-IL15 mice 2 h post HB-230 administration (HB-230 shown in magenta; CD11c shown in green; nuclei shown in blue). (B, C) WT and DdVil-IL15 mice were orally administered 10mg/kg HB-230 for 2 h before sacrifice, followed by staining with antibodies. The frequency of CD11c⁺MHCII⁺F4/80⁺ macrophages, CD3⁺ T cells, and CD19⁺ B cells was quantified among total CD45⁺ and HB-230⁺CD45⁺ cells. (D, E) The dot plots show the percentage of HB230^+^ cells within CD103⁺XCR1⁺ cDC1 and CD103⁺CD11b⁺ cDC2 subsets. Statistical significance was determined using two-way ANOVA with Fisher’s least significant difference (LSD) test (B, C) or one-way ANOVA with Tukey’s multiple comparison test (D, E) *, *p* < 0.05; ****, *p* < 0.001. Data represent mean ± SEM.

**Supplemental Figure 7.**
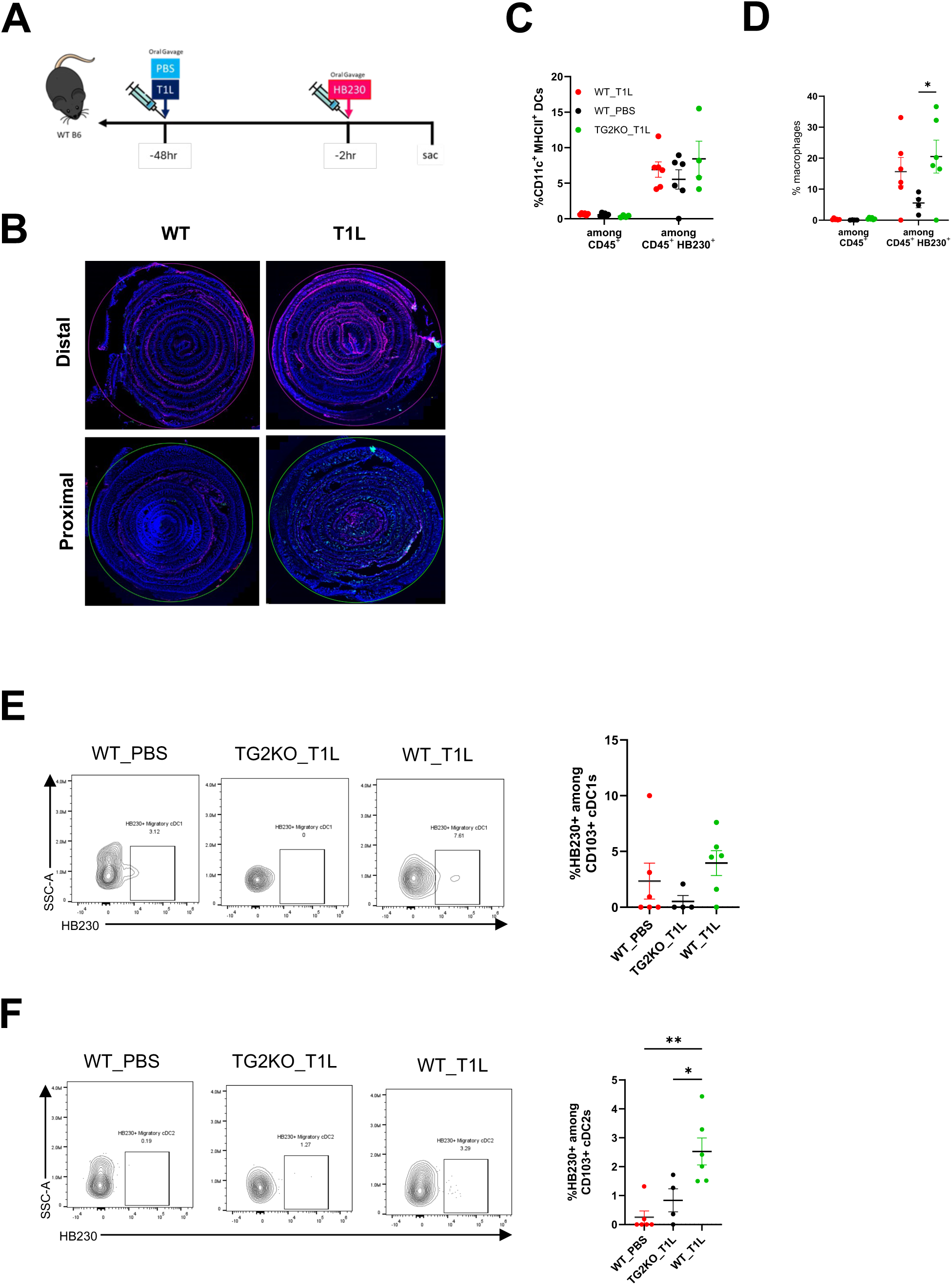
Reovirus infection increased ECM-associated TG2 activity and HB-230 uptake by CD103^+^CD11b^+^ cDC2. (A) Experimental schema of the reovirus infection model. WT and TG2KO mice were administered 3×10L PFU of T1L reovirus or PBS via oral gavage 48 h before receiving 10 mg/kg HB-230. Two hours following HB-230 dosing, mice were sacrificed, and the small intestines were collected for analysis. (B) Representative confocal images of distal and proximal Swiss-rolled small intestine sections (HB-230 shown in magenta; CD11c shown in green; nuclei shown in blue). Intestinal lamina propria cells were isolated from T1L-infected mice, followed by staining with antibodies. The dot plot indicates the percentage of CD11c⁺MHCII⁺F4/80⁻ DCs (C) and CD11c⁺MHCII⁺F4/80⁺ macrophages (D) among total CD45⁺ or HB-230⁺CD45⁺ cells. CD103^+^ DCs are subcategorized and analyzed for the percentage of HB230^+^ among CD103^+^XCR1^+^ cDC1 (E) or CD103^+^CD11b^+^ cDC2 (F). Statistical significance was determined using two-way ANOVA with Turkey’s multiple comparison test. (C, D) or one-way ANOVA with Tukey’s multiple comparison test (E, F). *, *p* < 0.05; **, *p* < 0.01. Data represent mean ± SEM.

**Supplemental Figure 8.**
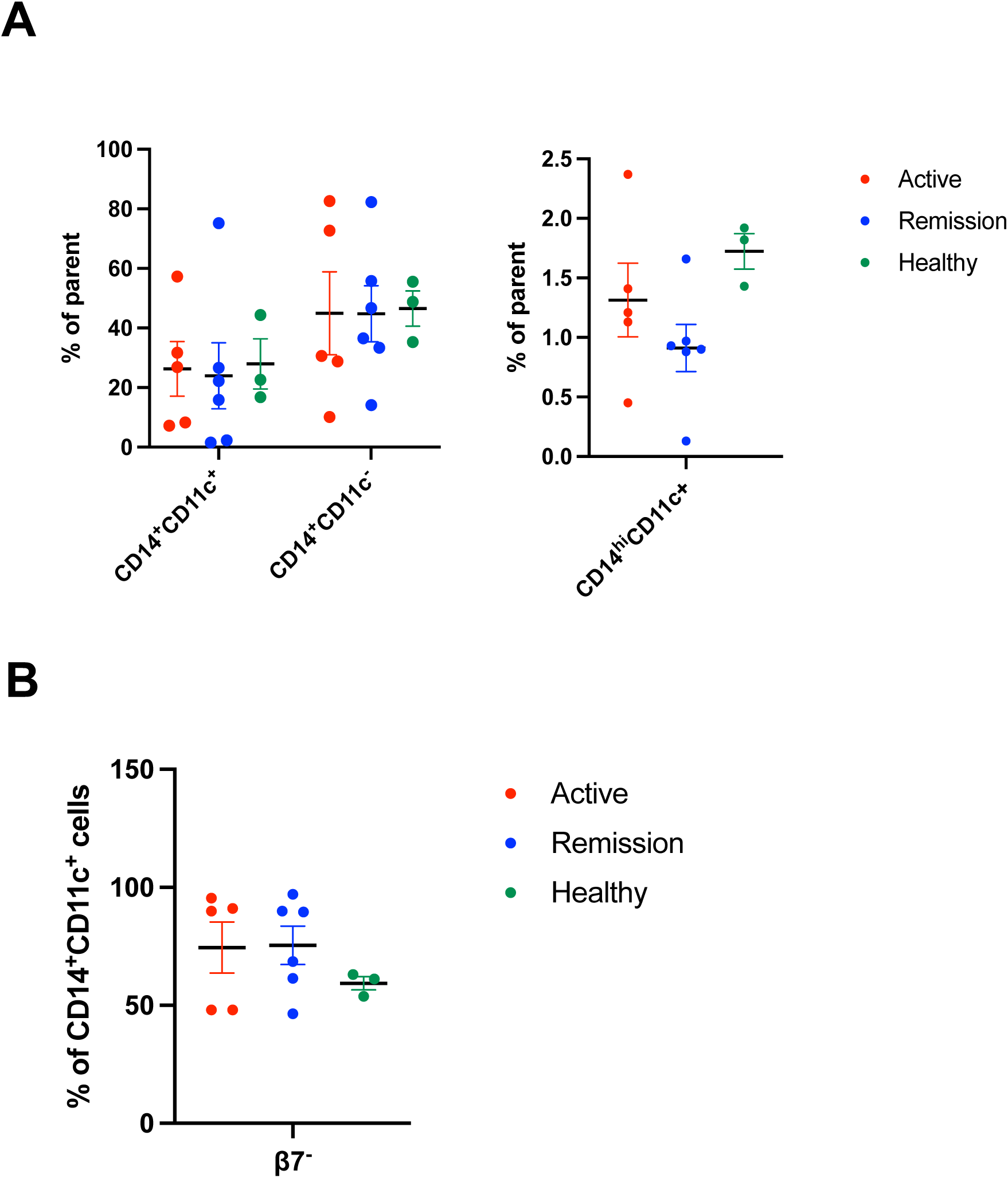
No significant differences are observed in the subpopulations of peripheral CD14^+^ monocytes between CeD patients and healthy subjects. Whole blood from either CeD patients or healthy subjects was collected, red blood cells were removed, and the remaining cells were incubated with HB-230. The cells were subsequently stained with antibodies and analyzed using a flow cytometer. (A) The dot plot displays the percentage of CD14^+^CD11c^+^, CD14^+^CD11c^-^, and CD14^hi^CD11c^+^ monocytes from CeD subjects or healthy donors. (B) The dot plot displays the percentage of β7^+^ cells among CD14^+^CD11c^+^ monocytes from CeD subjects (active, n=5; remission, n=6) or healthy donors (n=3). Data represent mean ± SEM.

